# Neural stemness unifies cell tumorigenicity and pluripotent differentiation potential

**DOI:** 10.1101/2021.12.23.474069

**Authors:** Min Zhang, Yang Liu, Lihua Shi, Lei Fang, Liyang Xu, Ying Cao

**Author notes:** Correspondence: Ying Cao, Model Animal Research Center of Medical School, Nanjing University, 12 Xuefu Road, Pukou High-Tech Zone, Nanjing 210061, China.

## Abstract

Tumorigenicity and pluripotent differentiation potential are kernel cell properties for tumorgenesis and embryogenesis. A growing number of studies have demonstrated that neural stemness is the source of the two cell properties, because neural stem cells and cancer cells share cell features and regulatory networks and neural stemness has an evolutionary advantage. However, it needs to validate whether neural stemness is a cell property that would unify tumorigenicity and pluripotent differentiation potential. SETDB1/Setdb1 is an epigenetic factor that is upregulated in cancer cells and promotes cancers, and correspondingly, is enriched in embryonic neural cells during vertebrate embryogenesis. We show that knockdown of SETDB1/Setdb1 led to neuronal differentiation in neural stem and cancer cells, concomitant with reduced tumorigenicity and pluripotent differentiation potential in these cells; whereas overexpression caused an opposite effect. On one hand, SETDB1 maintains a regulatory network comprised of proteins involved in developmental programs and basic cellular functional machineries, including epigenetic modifications (EZH2), ribosome biogenesis (RPS3), translation initiation (EIF4G), spliceosome assembly (SF3B1), etc., all of which play active roles in cancers. On the other, it represses transcription of genes promoting differentiation and cell cycle and growth arrest. Moreover, neural stemness, tumorigenicity and pluripotent differentiation potential were simultaneously enhanced during serial transplantation of cancer cells. Expression of proteins involved in developmental programs and basic cellular functional machineries, including SETDB1 and other proteins above, was gradually increased. In agreement with increased expression of spliceosome proteins, alternative splicing events also increased in tumor cells derived from later transplantations, suggesting that different machineries should work concertedly to match the status of high proliferation and pluripotent differentiation potential. The study presents the evidence that neural stemness unifies tumorigenicity and differentiation potential. Tumorigenesis represents a process of gradual loss of original cell identity and gain of neural stemness in somatic cells, which might be a distorted replay of neural induction during normal embryogenesis.

## Introduction

Neural stemness has usually been considered to be a type of tissue stemness. However, it exhibits some unique properties that are not displayed by other types of tissue stem cells. Neural stem cells (NSCs) exhibit tumorigenicity, because NSCs are capable of tumor formation when transplanted into immunodeficient mice (Cao, 2021; Xu et al., 2021), similar to cancer cells. NSCs have pluripotent differentiation potential, which means that, in addition to the capability of differentiation into cell types of the nervous system, they can also be induced to differentiate into different non-neural cell types (Cao, 2021; Clarke et al., 2000; Tropepe et al., 2001; Xu et al., 2021). Although embryonic stem cells (ESCs) have both tumorigenic and pluripotent differentiation potential, their intrinsic property is neural stemness because their default fate is primitive NSCs according to the “neural default model” of embryonic pluripotent cells (Cao, 2021; Gilbert and Barresi, 2016; Muñoz-Sanjuán and Brivanlou, 2002; Smukler et al., 2006; Tropepe et al., 2001). In somatic cells, blocking endogenous factors also leads to the gain of neural stemness and tumorigenicity at the expense of original cell identity (Cheng et al., 2014; Li et al., 2020; Southall et al., 2014; Xu et al., 2021). The association between neural stemness, tumorigenicity and pluripotent differentiation potential is supported by that most genes promoting tumorigenesis and pluripotency are neural stemness genes or genes with localized or at least enriched expression in embryonic neural cells (Cao, 2021; Zhang et al., 2017). The unique significance of neural stemness should be derived from the evolutionary advantage of neural genes and the neural-biased state in the last common unicellular ancestor of metazoans (Cao, 2021; Domazet-Lošo et al., 2007; Xu et al., 2021). We have suggested that neural stemness represents the ground state of tumorigenicity and pluripotent differentiation potential, and tumorigenesis is a process of gradual loss of original cell identity and restoration of neural ground state in somatic cells (Cao, 2017; Cao, 2021).

Neural ground state is also reflected by that genes involved in basic physiological functions in cells, such as cell cycle, ribosome biogenesis, proteasome or spliceosome assembly, protein translation, etc., are mostly enriched in neural cells during vertebrate embryogenesis. The genes involved in the basic machineries have usually a unicellular origin during evolution (Cao, 2021; Xu et al, 2021). Cell cycle is tightly linked to cell fate decision. Pluripotent stem cells or NSCs have a shorter cell cycle and differentiated cells have a longer one (Dalton, 2015), in agreement with the differential expression of cell cycle genes in neural and non-neural cells during embryogenesis (Cao, 2021). Cells with a fast cell cycle need a high protein production and homeostasis. Accordingly, genes for ribosome and proteasome proteins show a dominant expression in embryonic neural cells during embryogenesis (Cao, 2021), suggesting a high activity of protein synthesis and turnover in embryonic neural cells. Recently, we showed that this is concertedly regulated and required for the maintenance of neural stemness in both NSCs and cancer cells (Chen et al., 2021). Spliceosome is responsible for alternative splicing, a mechanism contributing to phenotypic novelty during evolution and to cell differentiation, lineage determination, tissue or organ formation during embryogenesis (Agosto and Lynch, 2018; Baralle et al., 2017; Bush et al., 2017). Enriched expression of genes involved in spliceosome assembly and hence alternative splicing matches the enrichment of long genes comprising of more exons/introns in both embryonic neural cells and cancer cells (Cao, 2021; Sahakyan and Balasubramanian, 2016; Xu et al., 2021). Moreover, genes involved in developmental programs, such as those for epigenetic modifications, pluripotency or promotion of pluripotency, etc., are also neural stemness genes or genes with specific or enriched expression in embryonic neural cells (Cao, 2021 and references therein). These genes work concertedly together to define a cellular state with a high proliferation and pluripotent differentiation potential, both being the features of tumorigenic cells. SETDB1 is an epigenetic modification factor responsible for trimethylation of lysine 9 of histone H3 (H3K9me3), and mediates transcriptional repression. It is required for early embryogenesis because gene knockout in mouse causes embryonic lethality before E7.5 (Dodge et al., 2004), suggesting that it’s critical during embryogenesis. Its gene is primarily transcribed in neural precursor cells during vertebrate embryogenesis, and brain-specific gene ablation causes severe brain developmental defect (Tan et al., 2012; Zhang et al., 2017). Accordingly, SETDB1 is upregulated in most cancers and promotes their initiation and progression (Strepkos et al., 2021). In the present study, by disruption of neural regulatory network via alteration of SETDB1/Setdb1 activity in cancer cells and NSCs and serial transplantation of cancer cells, we show that neural stemness is a cell property unifying tumorigenicity and pluripotent differentiation potential.

## Results

### Setdb1 is required for maintenance of neural stemness and differentiation potential of NSCs

Mouse ESCs turn into primitive NSCs (primNSCs) when cultured in defined NSC-specific serum-free medium (Ying et al., 2003; Smukler et al., 2006), which formed free-floating neurospheres in the medium (Figure 1A). Treatment of the neuropheres with retinoic acid (RA), a reagent inducing neuronal differentiation in NSCs, caused a neuronal differentiation phenotype (Figure 1A). RA treatment led to a decrease in neural stemness markers Sox1 and Msi1 in differentiated cells. Meanwhile, Myc, Hes1, Fgfr1, Vim and Cdh2, which are expressed specifically or dominantly in embryonic neural cells during vertebrate embryogenesis, and promote or are upregulated during tumorigenesis, were reduced in response to RA treatment (Figure 1B). By contrast, neuronal markers Map2 and Tubb3 are upregulated, supporting the neuronal differentiation effect (Figure 1B). We observed a decreased expression of epigenetic modification factors Setdb1 and Ezh2 in cells treated with RA (Figure 1B). Ezh2 mediates trimethylation of lysine 27 of histone H3 (H3K27me3) and hence transcription repression, and is required for early embryogenesis, maintaining or promoting neural stemness (Buschendorf et al., 2008; Lei et al., 2019; Sher et al., 2011). Intriguingly, proteins involved in translation initiation (Eif4g), ribosome biogenesis (Rps3 and Rpl26), and alternative splicing (Srsf3 and Sf3b1), were simultaneously reduced (Figure 1B). The result suggests that loss of neural stemness leads to a simultaneously decreased level of the machineries for basic cellular physiological functions and developmental programs.

**Figure 1.**
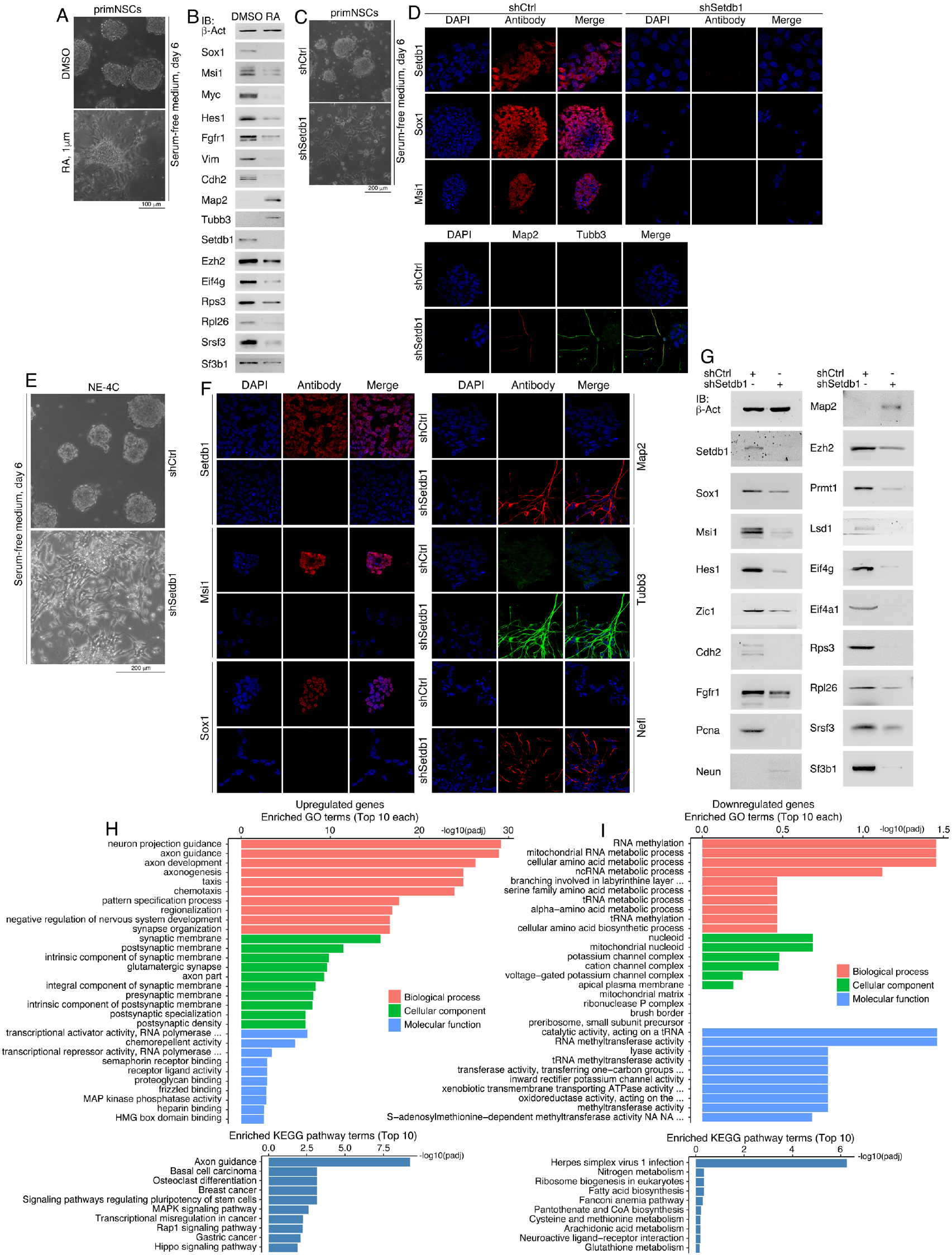
Neuronal differentiation of primNSCs induced by RA treatment and by knockdown of Setdb1. (A, B) Neuronal differentiation phenotype in ESC-derived primNSCs that were treated with RA at an indicated dose and cultured in NSC-specific serum-free medium for a period as indicated (A), and IB detection of protein expression in control (DMSO) and treated (RA) cells (B). (C, D) Neuronal differentiation phenotype in ESC-derived primNSCs induced by knockdown of Setdb1 and cultured in serum-free medium (C), and IF detection of protein expression in control (shCtrl) and knockdown (shSetdb1) cells (D). (E-G) Neuronal differentiation phenotype in NE-4C cells induced by knockdown of Setdb1 and cultured in serum-free medium (E), and IF (F) and IB (G) detection of protein expression in control (shCtrl) and knockdown (shSetdb1) cells. (H, I) Enrichment analysis on GO and KEGG pathway terms for upregulated (H) and downregulated (I) genes identified in a transcriptome profiling on control and knockdown cells in (E). In (D) and (F), cell nuclei were counterstained with DAPI.

Neuronal differentiation is accompanied with downregulation of Setdb1, we next examined how it functions in NSCs. Control primNSCs infected with lentivirus generated from empty vector (shCtrl) formed neurospheres in serum-free medium, knockdown of Setdb1 with a validated short hairpin RNA (shSetdb1) caused a failure of neurosphere formation. The resulting cells assumed a neuronal differentiation phenotype (Figure 1C). The short hairpin RNA could efficiently block Setdb1 expression in cells. Correspondingly, immunofluorescence (IF) showed that Sox1 and Msi1 were repressed in response to Setdb1 knockdown, but Map2 and Tubb3 were stimulated (Figure 1D), suggesting an essential role of Setdb1 in maintaining stemness in primNSCs.

We next explored whether this holds true for a different NSCs. NE-4C cells, a NSC cell line derived from cerebral vesicles of mouse E9 embryos, formed neurospheres in serum-free medium. When Setdb1 was knocked down, the cells displayed neuronal differentiation effect (Figure 1E). IF revealed that Sox1 and Msi1 were repressed when Setdb1 was lost, whereas neuronal proteins Map2, Tubb3 and Nefl were strongly stimulated (Figure 1F). Detection of expression of more proteins demonstrated that besides the repression of Sox1 and Msi1, neural stemness proteins and proteins with their gene expression localized to or enriched in embryonic neural cells, Hes1, Zic1, Cdh2, Fgfr1 and Pcna, were also downregulated (Figure 1G). By contrast, Neun and Map2 were upregulated (Figure 1G). Setdb1 knockdown caused a decreased expression of epigenetic factors Ezh2, Prmt1, and Lsd1 (Figure 1G), which maintain neural stemness in NSCs and cancer cells or confer neural stemness to differentiated cells (Chen et al., 2021; Han et al., 2014; Hirano et al., 2016; Lei et al., 2019; Sher et al., 2011). Moreover, a decreased expression in proteins involved in translation initiation (Eif4g, Eif4a1), ribosome biogenesis (Rps3, Rpl26) and alternative splicing (Srsf3, Sf3b1) was also observed (Figure 1G).

A transcriptome assay showed that Setdb1 knockdown caused an upregulation of 2619 genes and downregulation of 2457 genes in NE-4C cells (Table S1). The upregulated genes are primarily involved in events associated with neuronal differentiation (Figure 1H). Notably, among the upregulated genes were the *Hox* genes, which are critical for patterning of anterior-posterior body plan and specification of segment identity of tissues, including neural tissue, during embryogenesis. Some genes that are involved in cell cycle or growth arrest (*Bcl6*, *Cdkn1a*, *Eif2ak3*, etc.), DNA repair and apoptosis (*Gadd45a*, *Gadd45b*, *Gadd45g*, etc.) were also upregulated (Table S1). Downregulated genes are associated with RNA and amino acid metabolism (Figure 1I). This change might be related with the decrease in proteins involved in alternative splicing, ribosome biogenesis and protein translation in knockdown cells. Among downregulated genes include *Aspm*, *Cenpk*, *Enc1*, *Lin28a*, *Mcm8*, *Nucks1*, *Pou5f1*, *Utf1*, etc. (Table S1), which regulate pluripotency and neural stemness, transcription, cell cycle, etc. Taken together, inhibition of Setdb1 in NSCs caused neuronal differentiation and corresponding change in regulatory networks.

Subsequently, we observed that Setdb1 knockdown strongly inhibited cell invasion and migration (Figure S1A), and the capability of colony formation (Figure S1B, C). NSCs, such as NE-4C, are capable of forming xenograft tumors that contain cell types of three germ layers (Xu et al., 2021). As reported, NE-4C cells formed xenograft tumors in all six immunodeficient nude mice; whereas knockdown cells formed only smaller tumors in two out of six mice (Figure 2A-C; Table S2), an indication of lower tumorigenicity of knockdown cells. Expression of neural stemness genes or genes with enriched expression in embryonic neural cells during vertebrate embryogenesis was much lower in xenograft tumors formed by knockdown cells than in those formed by control cells (Figure 2D). The same tendency of difference in expression was observed for genes representing neuronal differentiation (Figure 2E) and genes representing mesodermal or endodermal tissue differentiation (Figure 2F). Immunohistochemistry (IHC) displayed that the expression level of Setdb1 was lower in sections of the tumors of knockdown cells than in tumors of control cells (Figure 2G), supporting the knockdown effect of Setdb1. Accordingly, neural stemness markers Sox1, Sox9, Pax6, and the proliferation marker Ki67, showed much weaker expression in tumors of knockdown cells (Figure 2G). In tumors of control cells, there was significant neuronal differentiation, as shown by Map2 expression; however, it was barely detectable in tumors of knockdown cells (Figure 2G). Similarly, markers for mesodermal and endodermal tissue differentiation, Acta2, Bglap, Ctsk, and Afp, were detected strongly in tumors of control cells, but detected weakly or not detected in tumors of knockdown cells (Figure 2G). These results demonstrate that loss of neural stemness in NE-4C cells via Setdb1 knockdown is accompanied by reduced potential of both tumorigenicity and pluripotent differentiation.

**Figure 2.**
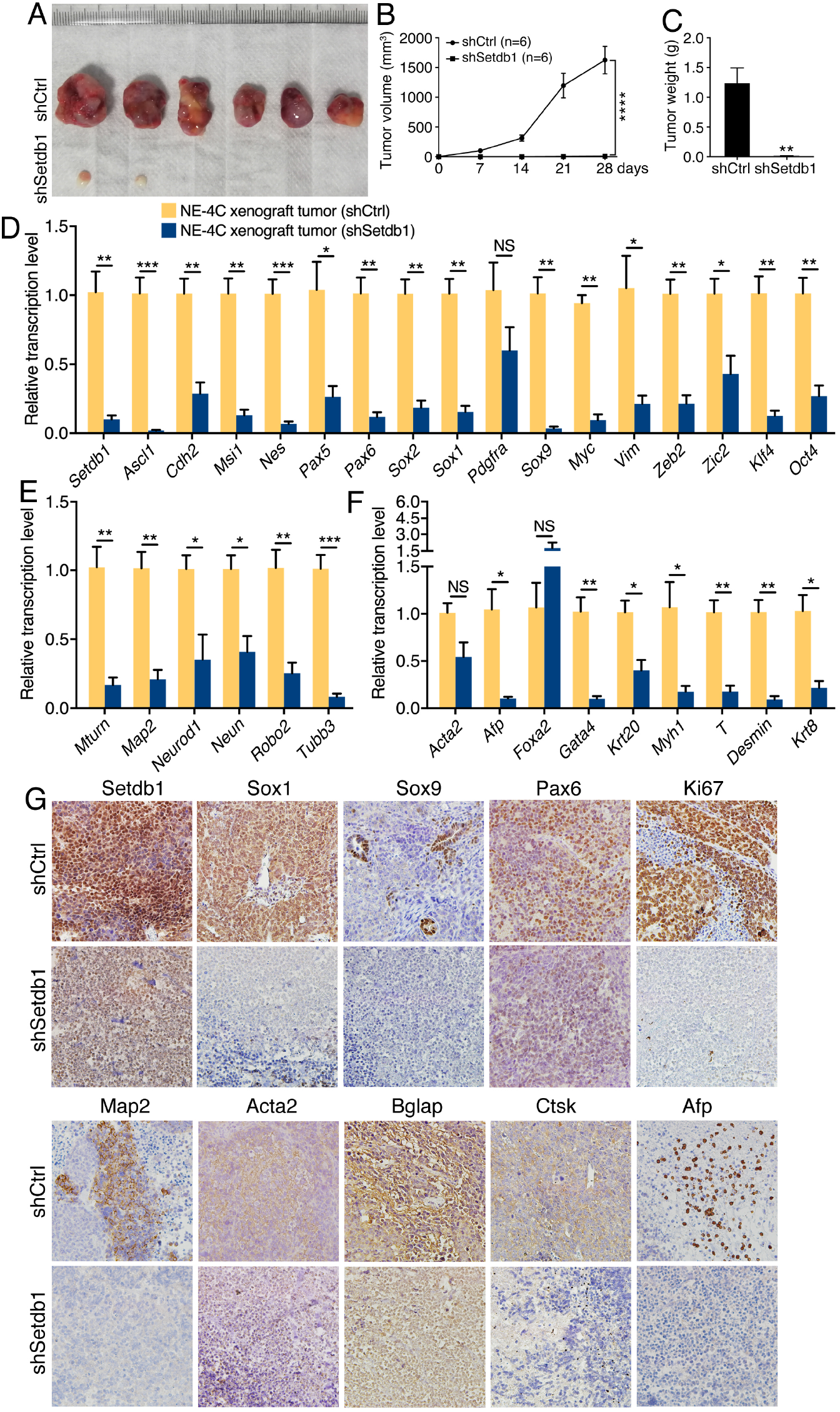
Difference in xenograft tumor formation and differentiation potential between control and knockdown NE-4C cells. (A-C) tumor formation of control (shCtrl) and knockdown (shSetdb1) cells in each six injected mice (A), and difference in tumor volume (B) and weight (C) between the two groups. Significance of difference in tumor volume (B) between two groups of mice was calculated using two-way ANOVA-Bonferroni/Dunn test. Significance of difference in tumor weight (C) was calculated using unpaired Student’s *t*-test. Data are shown as mean ± SEM. **p < 0.01, ****p < 0.0001. (D-F) RT-qPCR detection of differential expression of genes representing neural stemness or enriched in embryonic neural cells (D), neuronal differentiation (E), and mesodermal and endodermal tissue differentiation (F) between tumors of control NE-4C cells and of knockdown cells. Significance in transcription change was calculated based on experiments in triplicate using unpaired Student’s *t*-test. Data are shown as mean ± SEM. *p < 0.05, **p < 0.01, ***p < 0.001. NS: not significant. (G) IHC detection of cell/tissue markers in sections of tumors derived from control and knockdown cells. Objective magnification: 20×.

### SETDB1 knockdown in cancer cells leads to reduced neural stemness, tumorigenicity and pluripotent differentiation potential

We then investigated whether SETDB1 loss would also cause a similar effect in cancer cells. The colorectal cancer cell line HCT116 exhibits neural stemness because it expresses neural stemness markers, forms neurosphere-like structures in NSC-specific serum-free medium, and has pluripotent differentiation potential (Xu et al., 2021). In normal culture medium, knockdown of SETDB1 using a validated short-hairpin RNA (shSETDB1) led to a significant phenotypic change in HCT116 cells. The cells grew long neurite-like processes, suggesting an effect of neuronal differentiation (Figure 3A). When cultured in serum-free medium, control cells formed free-floating spherical structures, similar to NSCs. However, knockdown cells could not form such structures efficiently (Figure 3A), an indication of reduced neural stemness. Immunoblotting (IB) showed an efficient SETDB1 knockdown in cells, which was accompanied with a reduction in the level of H3K9me3. The cells express a series of neural stemness markers and proteins that are primarily expressed in embryonic neural cells during embryogenesis, including SOX2, MSI1, HES1, ZIC1, MYC, SOX9, OCT4, PCNA, CCND1, FGFR1, CDH2, the epigenetic factors EZH2, PRMT1, and LSD1, which promote cancers or are upregulated in cancer cells. They were all repressed in knockdown cells (Figure 3B). Reduced expression of CCND1 and PCNA indicates reduced proliferation in cells. Additionally, EIF4G, EIF4A1, RPS3, RPL26, SRSF3 and SF3B1, were downregulated (Figure 3B). Upregulated proteins were MAP2, NEUN and SYN1 (Figure 3B). IF also detected a reduced expression of SF3B1 and RPS3 but augmented expression of neuronal proteins NEUROD1, SYN1 and TUBB3 in knockdown cells (Figure 3C). Transcriptome profiling revealed that blocking SETDB1 in HCT116 cells caused downregulated transcription of genes that are significantly linked with biological processes of DNA replication, cell cycle, and corresponding cellular components, cellular functions and pathways (Figure S2A; Table S3). The upregulated genes are associated with endoplasmic reticulum to Golgi vesicle-mediated transport and processes of unfolded and topologically incorrect proteins (Figure S2B; Table S3). The upregulated genes in NE-4C knockdown cells, such as the *HOX* genes (*HOXA1*, *HOXB9*), *BCL6*, *CDKN1A*, *EIF2AK3*, *GADD45A*, *GADD45B*, *GADD45G*, etc., were also upregulated, and downregulated genes *ASPM*, *CENPK*, *ENC1*, *MCM8*, *NUCKS1*, etc., were downregulated in HCT116 knockdown cells (Table S3). Therefore, SETDB1 blocking in HCT116 caused a concerted decrease in basic cellular functional machineries (e.g., ribosome biogenesis, spliceosome assembly, translation), proteins involved in developmental programs, gene regulatory networks of cell proliferation, and a phenotype of neuron-like differentiation.

**Figure 3.**
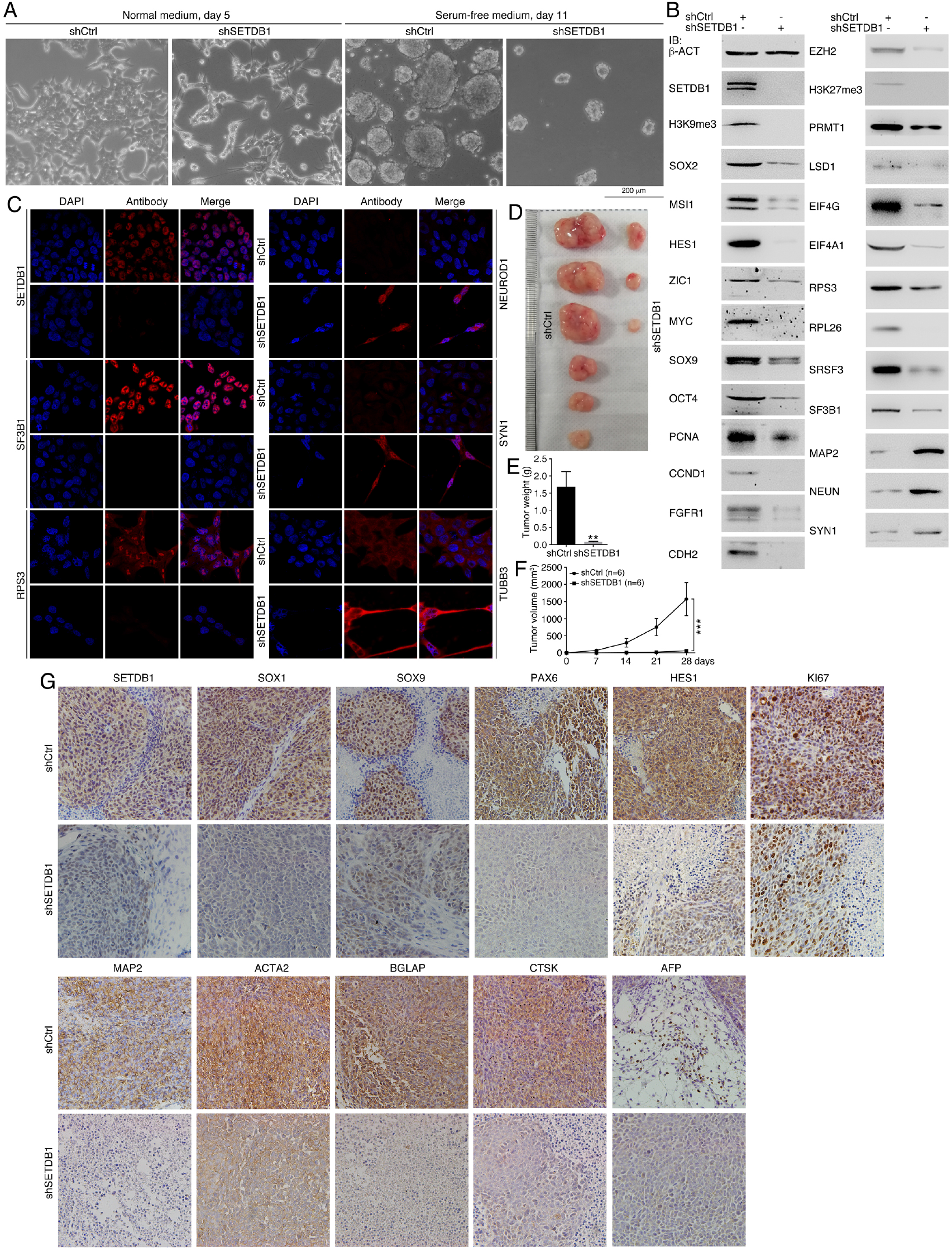
Influence of SETDB1 knockdown on the property of HCT116 cells. (A) Phenotypic change in HCT116 cells after SETDB1 knockdown and cultured in normal and serum-free medium in a period as indicated. (B, C) IB (B) and IF (C) detection of protein expression in control and knockdown cells. In (C), cell nuclei were counterstained with DAPI. (D-F) tumor formation of control and knockdown cells in each six injected mice (D), and difference in tumor weight (E) and volume (F) between the two groups. Significance of difference in tumor weight (E) was calculated using unpaired Student’s *t*-test. Significance of difference in tumor volume (F) was calculated using two-way ANOVA-Bonferroni/Dunn test. Data are shown as mean ± SEM. **p < 0.01, ***p < 0.001. (G) IHC detection of cell/tissue markers in sections of tumors derived from control and knockdown cells. Objective magnification: 20×.

Accordingly, the capability of invasion, migration and colony formation was compromised in HCT116 knockdown cells (Figure S2C-E). These cells showed much weaker ability of xenograft tumor formation (Figure 3D-F; Table S2). Tumors derived from control cells contained cells with high expression of SOX1, SOX9, PAX6 and HES1, and the proliferation marker KI67. Whereas in tumors of knockdown cells, they displayed weaker expression (Figure 3G), suggesting that these tumors contained cells with weaker neural stemness and proliferation. Control tumors had cells showing significant expression of MAP2, ACTA2, BDLAP or CTSK, the evidence of neuronal differentiation and mesodermal tissue differentiation. Detection of AFP in scattered cells means endodermal tissue differentiation. Tumors from knockdown cells exhibited less significant expression of these markers (Figure 3G). In summary, knockdown of SETDB1 in HCT116 cells led to reduced neural stemness, and consequently compromised tumorigenicity and differentiation potential, similar to the effect observed in NE-4C cells.

### Enhanced neural stemness by SETDB1 overexpression is accompanied with increased tumorigenicity and pluripotent differentiation potential

The colorectal cell line SW480 has a weaker tumorigenicity than cells such as HCT116. Correspondingly, SW480 couldn’t form spherical structures in serum-free medium, but HCT116 formed neurospheres within a same period of culture (Xu et al., 2021; Figure S3). We investigated how SETDB1 overexpression could affect the properties of SW480 cells. Overexpression of SETDB1 caused only weak change in cell morphology in normal culture medium. In serum-free medium, control cells formed clusters attached to the bottom of culture dish. Nevertheless, cells with overexpression formed free-floating spherical structures (Figure 4A), similar to NSCs or HCT116 cells. A series of neural stemness markers and proteins maintaining neural stemness, SOX1, MSI1, MYC, HES1, SOX9, FGFR1, and EZH2, were upregulated in overexpression cells (Figure 4B). IF also detected an enhanced expression of SOX1 and MSI1 in overexpression cells (Figure S4A). Increased CCND1 expression indicated that overexpressed SETDB1 promoted cell cycle. Contrary to SETDB1 knockdown, overexpression enhanced expression of EIF4G, RPS3, RPL26, SRSF3, and SF3B1 (Figure 4B). This reflects again that basic cellular functional machineries should be concertedly regulated to match the proliferation rate of cells, as reported previously (Cao, 2021; Chen et al., 2021). Repressed expression was observed for the epithelial marker CDH1 and for tumor suppressors RB1 and TP53 (Figure 4B), which functions primarily in cell cycle arrest, maintaining genomic integrity and cell differentiation, etc.

**Figure 4.**
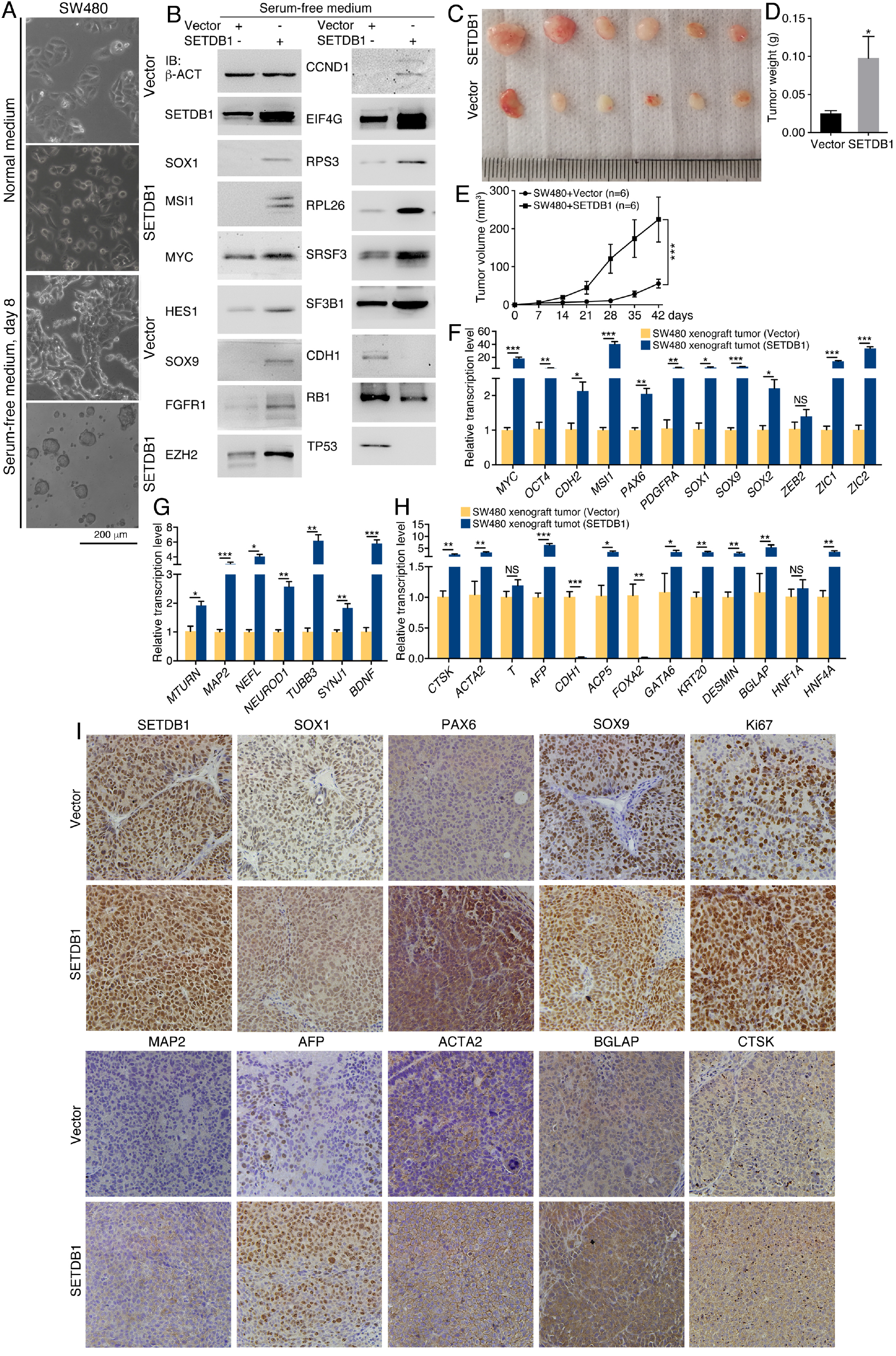
Influence of SETDB1 overexpression on the property of SW480 cells. (A) Comparison of phenotypic change between control (Vector) SW480 cells and cells with SETDB1 overexpression (SETDB1), which were cultured in normal and serum-free medium, respectively, in a period as indicated. (B) Protein expression change between control and overexpression cells, as detected with IB. (C-E) tumor formation of control and overexpression cells in each six injected mice (C), and difference in tumor weight (D) and volume (E) between the two groups. Significance of difference in tumor weight (D) was calculated using unpaired Student’s *t*-test. Significance of difference in tumor volume (E) was calculated using two-way ANOVA-Bonferroni/Dunn test. Data are shown as mean ± SEM. *p < 0.05, ***p < 0.001. (F-H) Difference in transcription of genes representing neural stemness (F), neuronal differentiation (G), and mesodermal and endodermal tissue differentiation (H) between tumors of control and of overexpression cells, as detected with RT-qPCR. Significance in transcription change was calculated based on experiments in triplicate using unpaired Student’s *t*-test. Data are shown as mean ± SEM. *p < 0.05, **p < 0.01, ***p < 0.001. NS: not significant. (I) IHC detection of cell/tissue markers in sections of tumors derived from control and overexpression cells. Objective magnification: 20×.

Enhancement of neural stemness in SW480 cells due to SETDB1 overexpression would mean an increment in tumorigenicity and differentiation potential. Indeed, we observed an increased capability of invasion, migration and colony formation (Figure S4B-D) in cells with SETDB1 overexpression. These cells displayed stronger ability of xenograft tumor formation than control cells (Figure 4C-E; Table S2). Detection of gene transcription demonstrated that a series of genes representing neural stemness or/and being involved in the maintenance of neural stemness were upregulated in xenograft tumors from overexpression cells (Figure 4F). Increased transcription was also detected for genes representing neuronal differentiation (Figure 4G) and genes representing mesodermal and endodermal cell differentiation (Figure 4H). IHC revealed a heavier staining for SETDB1 in sections of tumors derived from overexpression cells than that in sections from control cell tumors (Figure 4I). Stronger signals were also observed for SOX1, PAX6, SOX9 and KI67 (Figure 4I), indicating that tumors of overexpression cells contained cells with elevated property of neural stemness and proliferation. Moreover, MAP2 was barely detected in tumors of control cells, but detected in tumors of overexpression cells. AFP, ACTA2, BGLAP and CTSK were more strongly detected in sections of tumors of overexpression cells (Figure 4I). Taken together, these results suggest that cells with enhanced neural stemness have a stronger capability to form tumors, which contain cells with elevated property of neural stemness and proliferation, and cells characteristic of neuronal, mesodermal and endodermal differentiation.

### SETDB1 sustains a regulatory network maintaining neural stemness

Inhibition of SETDB1 results in a decrease in the expression of proteins involved in the functions of basic cellular functional machineries and developmental programs. These let us to investigate whether SETDB1 might regulate these proteins or their genes. Mass spectrometry identified 222 putative Setdb1 interaction partners in NE-4C cells (Table S4). They are mostly involved in pathways of ribosome and spliceosome, which are responsible for the biological processes of translation, mRNA processing and RNA splicing, etc. Correspondingly, these proteins are mainly associated with molecular functions and cellular components that are involved in such pathways and biological processes, such as RNA binding, ribosome constituents, intracellular ribonucleoprotein complex, cytoplasm, nucleus, etc. (Figure S5; Table S4). In HCT116 cells, SETDB1 was found to interact with EZH2, SF3B1, RPS3, RPL26 and EIF4G. SETDB1 also interacted with proteasome component proteins PSMD2 and ADRM1, and USP11 (Figure 5A), a deubiquitinase that is expressed primarily in embryonic nervous system, regulates neurogenesis and neuronal migration (Chiang et al., 2021), and accordingly, is upregulated in cancer cells and promotes cancers (Guo et al., 2019; Lazaro-Camp et al., 2021; Wang et al., 2019).

**Figure 5.**
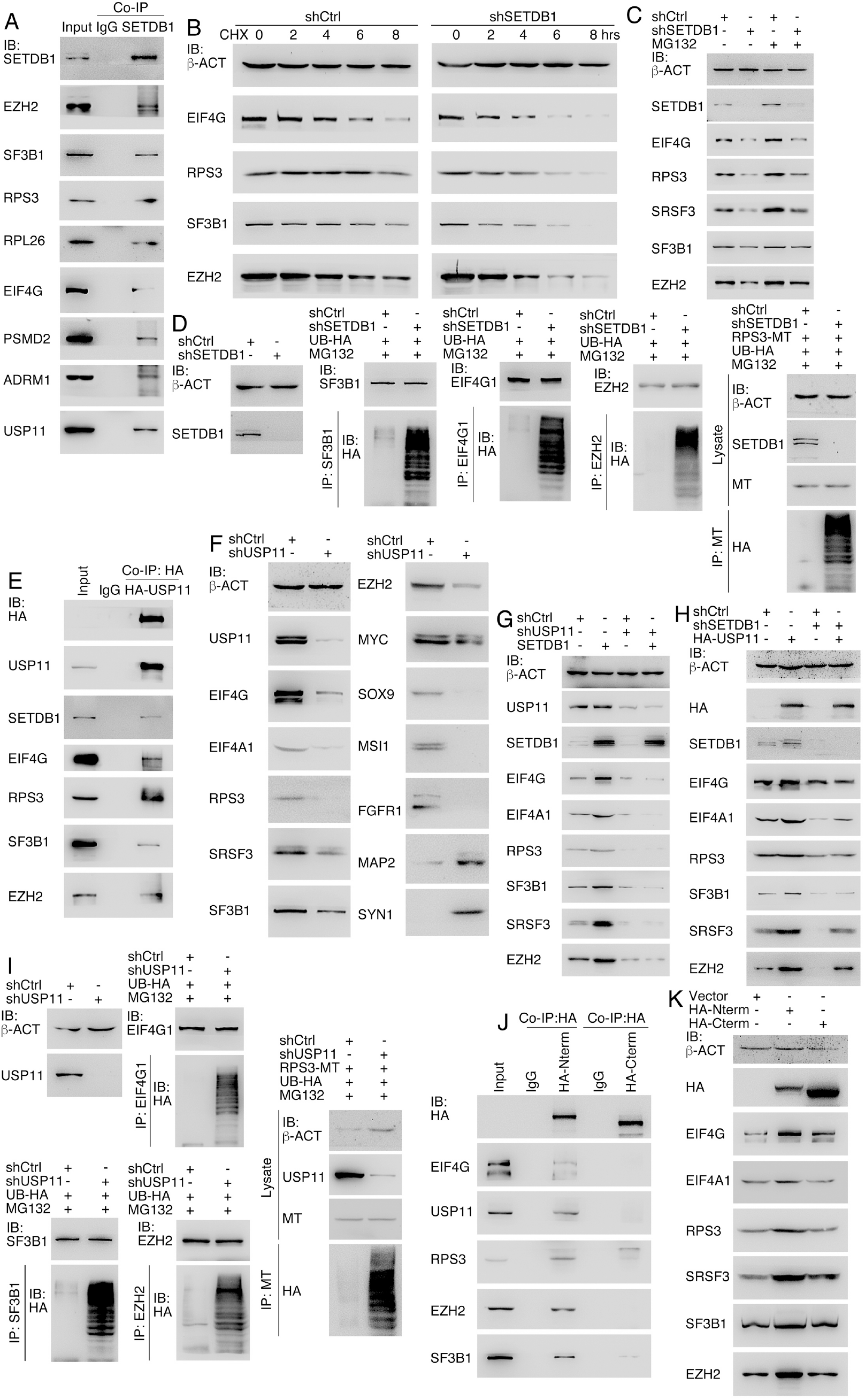
SETDB1 maintains protein stability via interaction with USP11. (A) Co-IP detection of binding of SETDB1 to other proteins. (B) The effect of protein degradation in response to inhibition of de novo protein synthesis via CHX treatment in a time series and SETDB1 knockdown in HCT116 cells. (C) Detection of the dependence of reduced protein expression caused by SETDB1 knockdown on proteasomal activity via treatment of cells with MG132. (D) Effect of SETDB1 knockdown on ubiquitination of endogenous SF3B1, EIF4G1, EZH2, and overexpressed RPS3, which were precipitated with their respective antibodies or the Myc-tag (MT) antibody. (E) Co-IP detection of binding of overexpressed HA-USP11 to other proteins. (F) Effect of USP11 knockdown on proteins expression in HCT116 cells. (G) Effect of USP11 knockdown on enhanced protein expression caused by overexpressed SETDB1. (H) Effect of SETDB1 knockdown on enhanced protein expression caused by overexpressed USP11. (I) Effect of USP11 knockdown on ubiquitination of endogenous EIF4G1, SF3B1, EZH2, and overexpressed RPS3, which were precipitated with their respective antibodies or the MT antibody. (J) Differential binding of N- and C-terminal region of SETDB1 to proteins. (K) Differential effect of enforced expression of N- and C-terminal region of SETDB1, respectively, on protein expression.

A time-course assay revealed that EIF4G, RPS3, SF3B1 and EZH2 degraded faster in HCT116 cells with SETDB1 knockdown than in control cells (Figure 5B). Since knockdown of SETDB1 did not cause a significant change in transcription of genes for these proteins (Figure S6A), we deduced that SETDB1 might regulate the stability of these proteins. Blocking SETDB1 reduced EIF4G, RPS3, SRSF3, SF3B1 and EZH2, while inhibition of proteasome activity with MG132 enhanced these proteins. Inhibition of proteasome activity was able to reverse the reduction of these proteins (Figure 5C), suggesting that SETDB1 mediated expression of these proteins depends on the ubiquitin-proteasome system. Indeed, SETDB1 knockdown caused an enhancement of ubiquitination of SF3B1, EIF4G1 and EZH2 that were precipitated with their respective antibodies, as well as overexpressed RPS3 that was precipitated with a Myc-tag (MT) antibody (Figure 5D). The ubiquitination status might be due to the interaction between SETDB1 and USP11. Co-IP showed that USP11 was able to interact with SETDB1, EIF4G, RPS3, SF3B1 and EZH2 (Figure 5E), indicating that USP11 and SETDB1 form complexes with other proteins. Similar to the effect of blocking SETDB1, knockdown of USP11 using a validated short hairpin RNA (shUSP11) in HCT116 cells led to a decreased level of EIF4G, EIF4A1, RPS3, SRSF3, SF3B1, EZH2, MYC, SOX9, MSI1, FGFR1, and increased level of MAP2 and SYN1 (Figure 5F). Although overexpression of SETDB1 enhanced protein expression, it could not do so in the absence of USP11 (Figure 5G). Contrarily, overexpression of USP11 caused enhanced expression of proteins and was able to rescue the reduced expression caused by SETDB1 knockdown (Figure 5H). Therefore, maintenance of protein expression by SETDB1 depends on USP11. Indeed, we found an enhanced ubiquitination of EIF4G1, SF3B1, EZH2 and RPS3 when USP11 was blocked (Figure 5I). We made two truncation mutants that contained the N-terminal region (aa 1-725) and C-terminal region (aa 726-1291) of SETDB1, and were fused with HA-tags, designated as HA-Nterm and HA-Cterm respectively. Forced expression of HA-Nterm revealed interaction with EIF4G, USP11, RPS3, EZH2 and SF3B1, whereas HA-Cterm did not (Figure 5J). In agreement, forced expression of HA-Nterm enhanced the expression of EIF4G, EIF4A1, RPS3, SRSF3, SF3B1 and EZH2, but forced expression of HA-Cterm did not display such an effect (Figure 5K). Since the pre-SET, SET and post-SET domains in the C-terminal region of SETDB1 are required for methyltransferase activity, differential effect of N- and C-terminal regions on protein interaction and expression suggests that the methyltransferase activity might be not essential for promoting protein stability.

SETDB1 regulates expression of EZH2, both mediating transcription silencing. Meanwhile, SETDB1 knockdown caused activation of genes promoting neuronal differentiation, cell cycle and growth arrest (Figure S6B). Accordingly, SETDB1 knockdown caused a decreased binding of EZH2 to the promoters, and correspondingly, a decreased level of H3K9me3 and H3K27me3 in the promoters of genes that are upregulated in NE-4C or/and HCT116 cells, e.g. *CDKN1A*, *GADD45B*, *BCL6*, *NEUROD1*, and *TUBB3* (Figure S6C). Decrease in these modifications change chromatin configurations into a state of transcriptional activation. In summary, these results suggest that SETDB1 functions in two layers. One is to bridge USP11 to the interaction partners, thereby preventing them from ubiquitination and degradation, the other is to establish a transcriptional silencing state in the promoters of genes promoting differentiation. As a result, SETDB1 maintains a high level of basic cellular functional machineries and factors involved in developmental programs to match tumorigenic and differentiation potentials in neural stem and cancer cells.

### Serial transplantation of SW480 cells demonstrate that neural stemness, tumorigencity and differentiation potential are coupled together

We analyzed whether neural stemness, tumorigencity and differentiation potential are coupled cell propertied using serial transplantation or in vivo passaging via subcutaneous injection in nude mice. The strategy was depicted in Figure S7A. Cells derived from xenograft tumors were cultured in NSC-specific serum-free medium for 9 days to test their ability to form neurosphere-like structures, an indication of neural stemness. In vitro cultured SW480 cells were designated as s0. Xenograft tumors and cells forming neurosphere-like structures from the first, second and the third passages in vivo were designated as s1, s2 and s3, respectively (Figure S7A). While SW480 cells did not form neurospheres in serum-free medium, cells from the first transplantation (s1) formed small spherical structures, whereas the cells from the second (s2) grew larger and those from the third (s3) grew the largest spherical structures (Figure 6A). Correspondingly, there was a tendency of increasing expression of MSI1 and SOX1 in cells or cell spheres from s0 to s3. The increasing expression was also observed for EIF4G, SF3B1, SRSF3 and RPS3 (Figure 6B). Similar expression tendency was observed for some additional proteins, including SETDB1 and its corresponding histone modification H3K9me3, SOX2, SOX1, MSI1, MYC, SOX9, CDH2, CCND1, PCNA, RPL26, EZH2 and H3K27me3 (Figure 6C). By contrast, CDH1, RB1 and TP53, which are repressed during tumorigenesis, were repressed during serial transplantation (Figure 6C). These data demonstrate that serial transplantation causes SW480 cells to enhance neural stemness gradually and the expression of proteins that promote neural stemness.

**Figure 6.**
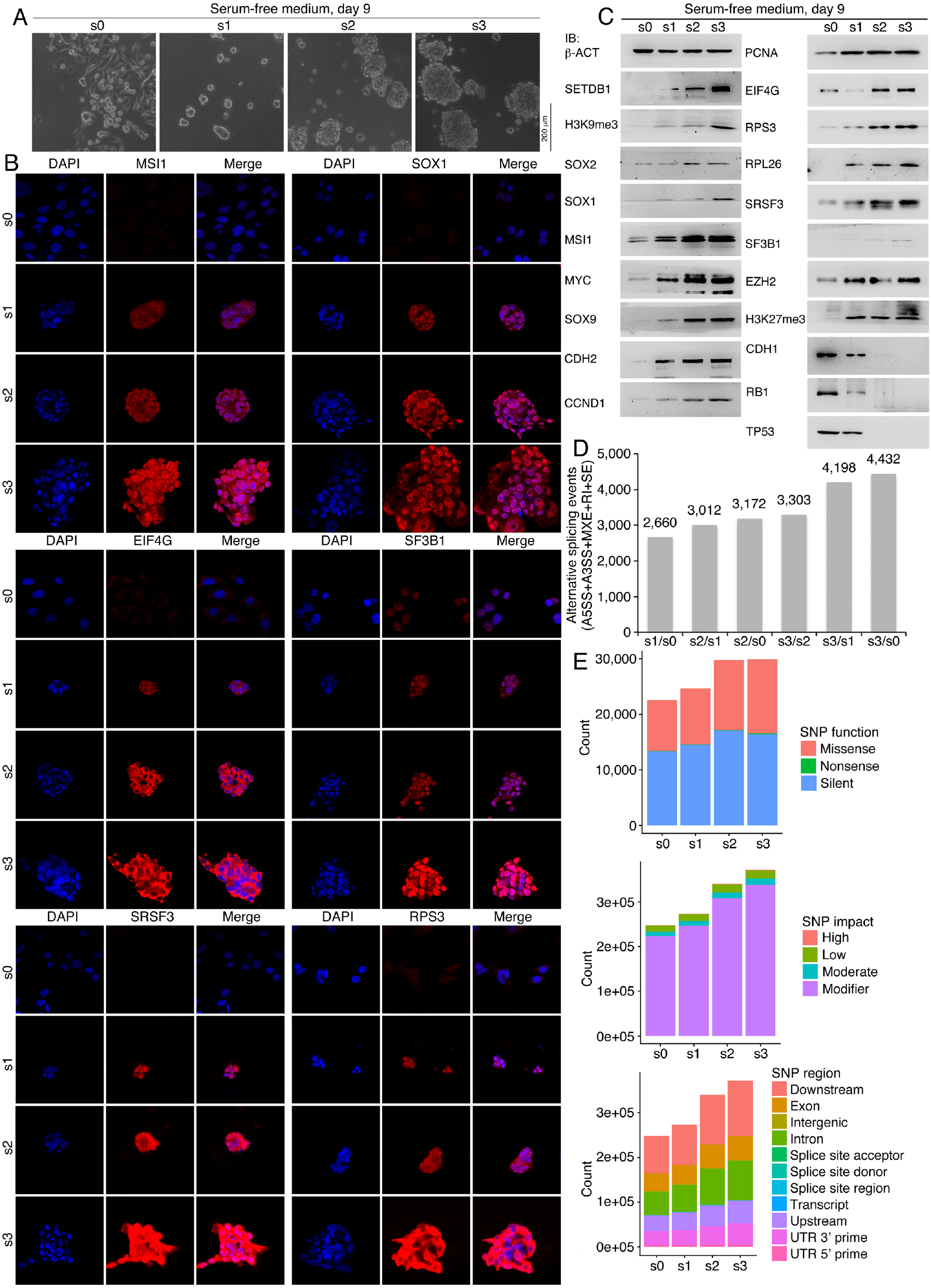
Characterization of property of SW480 cells and cells derived from tumors by serial transplantation. (A) Different ability of cell sphere formation in serum-free medium by SW480 cells (s0) and cells derived from tumors by serial transplantation (s1, s2 and s3). (B, C) Detection of protein expression with IF (B) and IB (C) in cells and cell spheres in (A). (D) Alternative splicing events compared between cells or cell spheres in (A) that were identified by RNA-sequencing. Abbreviations: A5SS, Alternative 5’ splice site; A3SS, Alternative 3’ splice site; MXE, Mutually exclusive exon; RI, Retained intron; SE, Skipped exon. (E) Numbers of SNPs in cells or cell spheres in (A) that were identified by RNA-sequencing, according to SNP function, impact or region of occurrence.

Overall transcription in cell spheres from s0 to s3 changed progressively. The transcriptome of s3 cell spheres displayed the biggest difference from that of s0 cells, with s2 cell spheres showing weaker difference and s1 spheres showing the weakest (Figure S7B). Interestingly, downregulated genes between s3 and s0, and s2 and s0 were strongly enriched in immune response and immune system process, suggesting that successive passaging in vivo leads to a gradually altered immune response in the cells (Figure S7C). This might reflect the growing ability of immune evasion of cancer cells during cancer progression (Gonzalez et al., 2018). Upregulated genes were weakly associated with cell adhesion (Figure S7D). The numbers of alternative splicing events were constantly increasing in cells with high passages as compared with cells with low passages (Figure 6D; Table S5). This corresponds with the increased expression of proteins involved in alternative splicing in cells from later passages. Meanwhile, the numbers of single nucleotide polymorphisms (SNPs) were also increasing during passaging (Figure 6E; Table S6), an effect associated with the decreased expression of TP53. The increase in SNPs and genomic abnormalities has been observed during cancer progression (Li et al., 2008; Wistuba and Meyerson, 2008).

The results above suggest that SW480 cells might have enhanced malignant features and tumorigenicity after passaging in vivo. Indeed, cells of passage 0, 1, 2 and 3 displayed constantly growing ability in invasion, migration (Figure S8A) and colony formation in soft agar (Figure S8B, C). During serial transplantation, cells formed small xenograft tumors in nude mice at the first time. Cells derived from the first transplantation formed larger tumors and tumors formed by cells from the second transplantation were even larger (Figure 7A-C; Table S2). Genes representing neural stemness or/and promoting cancers, *SETDB1*, *SOX1*, *MSI1*, *PAX6*, *SOX2*, *MYC* and *ZIC2* showed a tendency of enhanced transcription during serial transplantation (Figure 7D). Analysis on the sections of the tumors revealed a similar tendency of increased expression of SETDB1, SOX1, SOX9 and KI67, and signals for these proteins were more densely stained in sections of tumors at higher passages than those at lower passages (Figure 7E). Additionally, MAP2, BGLAP, CTSK and AFP were more strongly stained in tumors from later passages (Figure 7E), suggesting that these tumors showed more efficient differentiation of tissue or cell types. In summary, serial transplantation leads to progressive enhancement of neural stemness, tumorigenicity and differentiation potential in tumorigenic cells.

**Figure 7.**
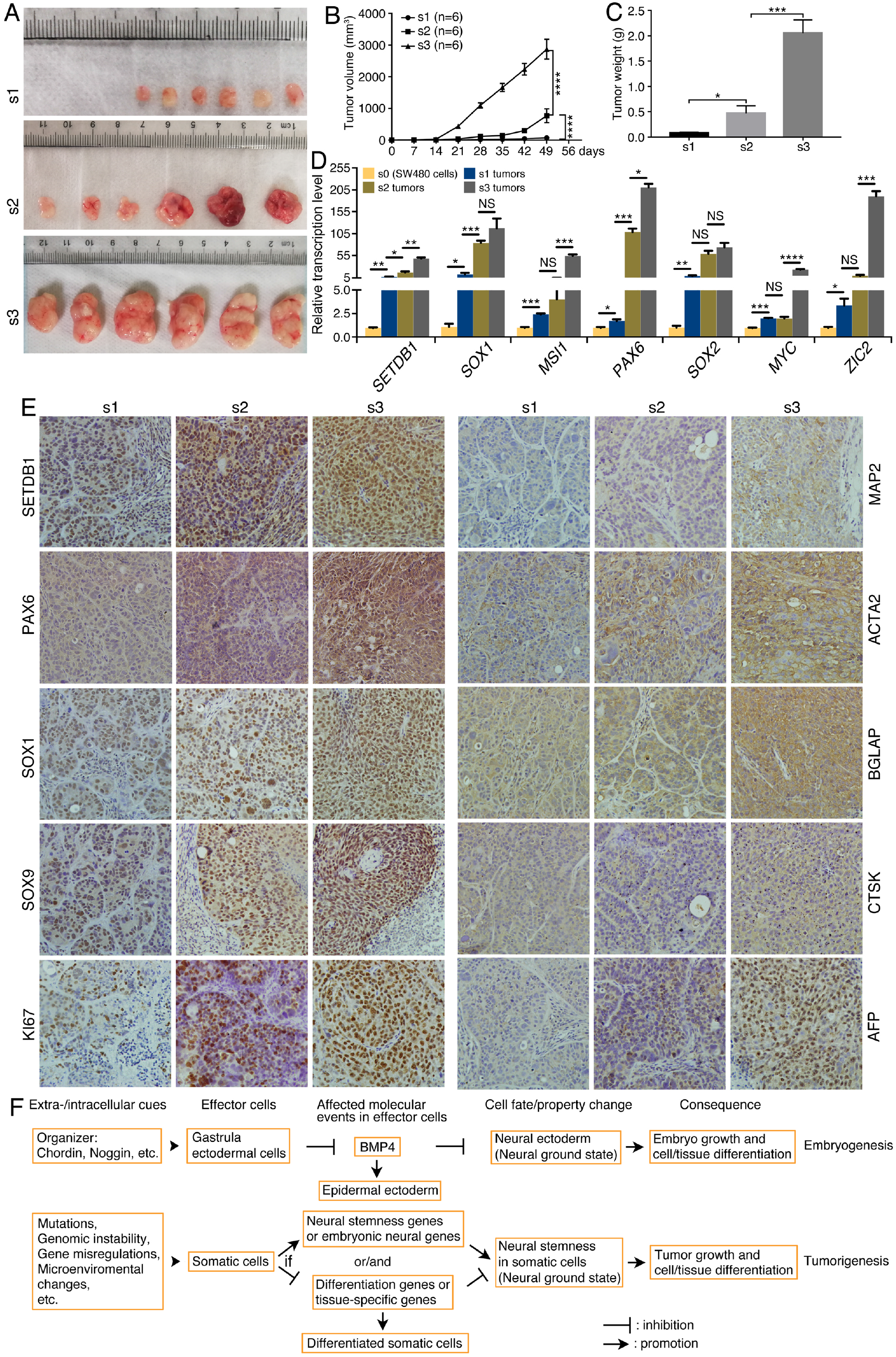
Characterization of xenograft tumors from SW480 cells by serial transplantiation and a general model unifying embryogenesis and tumorigenesis. (A-C) tumor formation via serial transplantation in each six injected mice (A), anddifference in tumor volume (B) and weight (C) between different groups. Significance of difference in tumor volume (B) was calculated using two-way ANOVA-Bonferroni/Dunn test. Significance of difference in tumor weight (C) was calculated using unpaired Student’s *t*-test. Data are shown as mean ± SEM. *p < 0.05, ***p < 0.001, ****p < 0.0001. (D) Comparison of expression of genes for SETDB1 and neural stemness protein in SW480 cells and xenograft tumors in (A), as revealed by RT-qPCR. Significance in transcription change was calculated based on experiments in triplicate using unpaired Student’s *t*-test. Data are shown as mean ± SEM. *p < 0.05, **p < 0.01, ***p < 0.001, ****p < 0.0001. NS: not significant. (E) IHC analysis of SETDB1 and other marker protein expression in sections of tumors in (A). Objective magnification: 20×. (F) A model depicting the correlation between embryonic development and tumorigenesis. Normal embryogenesis needs the inhibition of epidermal fate in gastrula ectoderm by extracellular signals secreted by Spemann organizer and hence restoration of the neural ground state in ectoderm. Formation of neural precursor tissues is required for differentiation of the nervous system and many non-neural tissues. This neural induction can occur ectopically during embryogenesis, caused by either an ectopic organizer activity or ectopic expression of embryonic neural genes, leading to formation of secondary embryonic structures. Nevertheless, this process might occur in any cells in any time of an animal life. Somatic cells could suffer various extracellular (e.g., microenvironmental change) or/and intracellular (e.g., mutations) insults. If occasionally the insults cause an activation of neural stemness regulatory network or/and downregulation/silencing of tissue-specific or differentiation genes/factors, then somatic cells lose their original cell identity and restore the neural ground state, similar to the case of gastrula ectodermal cells, and tumorigenesis initiates and progresses, resembling a defected process of embryonic development. For detailed information, see text and also refer to Cao (2017, 2021) and references therein.

## Discussion

Tumorigenicity and pluripotent differentiation potential are the basic cell properties for understanding tumorigenesis and embryogenesis, respectively. Although ideas like “cancer as a dynamic developmental disorder” were proposed previously (Rubin, 1985), the most prominent connection of cancer and embryonic development might be exemplified by teratocarcinoma. In recent few years, our studies generalize that cancer cells exhibit neural stemness and pluripotent differentiation potential. In combination with other studies, we suggest that neural stemness might be the ultimate determinant for these basic cell properties (Cao, 2017; Cao, 2021; Xu et al., 2021; Zhang et al., 2017). This notion is further supported by the present study. When the network regulating neural stemness was disrupted, for example, via inhibition of Setdb1/SETDB1, NSCs or cancer cells exhibited neuronal or neuronal-like differentiation, concomitant with reduced malignant features, tumorigenicity and differentiation potential in the cells. Vice versa, overexpression caused simultaneously enhanced neural stemness, tumorigenicity and differentiation potential in mildly tumorigenic cells. Importantly, these cell properties were getting stronger during the process of serial transplantation, which is believed to mimic cancer progression (Chan et al., 2007). These lines of evidence demonstrate that tumorigenicity and pluripotent differentiation potential are integral properties of neural stemness.

A cell property is controlled by a regulatory network rather than a single molecule or molecular event. Cancer cells and NSCs share a regulatory network, which is primarily comprised of factors with specific or enriched expression in NSCs or embryonic neural cells (Cao, 2017; Cao, 2021; Xu et al., 2021; Zhang et al., 2017). Enriched expression of SETDB1 in embryonic neural cells and its functions during neuronal differentiation and neural development means that it is a component of neural regulatory network and a component of regulatory network promoting cancers. By recruiting deubiquitinase USP11, SETDB1 interacts with and maintains expression of components in the regulatory network, including neural stemness proteins and proteins involved in translation initiation, ribosome biogenesis and spliceosome assembly, in cancer cells and NSCs. Potential interaction partners of SETDB1 also include proteins participating DNA replication and transcription (MCM3, TOP1, TOP2A, DDX3X) (Table S4), nucleocytoplasmic transport of proteins and RNAs (RAN), proteasome assembly (PSMD2, ADRM1), etc., suggesting that SETDB1 might regulate a broader range of proteins that are involved in basic cellular functional machineries, which are usually enriched in embryonic neural cells (Cao, 2021; Xu et al., 2021). Like that PRMT1 coordinates ribosome and proteasome to maintain neural stemness in cancer cells and NSCs (Chen et al., 2021), SETDB1 plays a role in the regulation of basic cellular functional machineries, thereby maintaining neural stemness, tumorigenicity and pluripotent differentiation potential in tumorigenic cells.

Genes regulating cancers and neural genes are both enriched in long genes consisting of more exons/introns as compared with non-neural genes (Cao, 2021; Sahakyan and Balasubramanian, 2016; Xu et al., 2021). Compatible with this feature is the enriched expression of components involved in spliceosome assembly in embryonic neural cells and cancer cells (Cao, 2021), suggesting a high activity of alternative splicing in these cells. Our data on serial transplantation of SW480 cells provides the convincing evidence for the association between spliceosome protein expression and alternative splicing. Cells from later passages, which exhibited stronger differentiation potential, displayed higher level of spliceosome protein expression. Accordingly, there were more alternative splicing events. It is logical that cells with enriched expression of long genes and high activity of alternative splicing are more plastic in differentiation potential. Generation of more splice variants means the need for more protein translation to execute their functions, and the need for more efficient protein turnover, etc. In general, all machineries should operate at a higher efficiency to match the status of fast cell cycle, high proliferation and pluripotent differentiation potential (Cao, 2021; Chen et al., 2021; present study). Thus, neural stemness should be the general stemness or the ground state for cell differentiation (Cao, 2021; Clarke et al., 2000). These basic cell functional machineries, such as cell cycle, ribosome, proteasome, spliceosome, epigenetic modifications, etc., have all be proved to play active roles during tumorigenesis. Inhibition of the activity of these machineries has been either applied to cancer therapy or under clinical trials (Bennett and Licht, 2018; Bhat et al., 2015; Bustelo and Dosil, 2018; Chen et al., 2017; Dvinge et al., 2016; Mohammad et al., 2019; Pelletier et al., 2018; Soave et al., 2017; Thibaudeau and Smith, 2019; Wang and Lee, 2018). Considering contribution of neural stemness to cell tumorigenicity and similarity in regulatory network between cancer and NSCs (Cao, 2017; Cao, 2021; Zhang et al., 2017), inhibition of cancer via targeted therapy is per se achieved by disruption of neural regulatory network.

NSCs are the obvious precursors of the nervous system. Contribution of neural stemness to non-neural differentiation is non-obvious. Others and we demonstrated the feature of pluripotency of NSCs in chimera embryo and xenograft tumor formation (Clarke et al., 2000; Tropepe et al., 2001; Xu et al., 2021). In fact, non-neural differentiation of embryonic neural cells can be observed during and is critical for normal embryogenesis. Locating between neural plate (the precursor tissue of central nervous system) and epidermal ectoderm, neural crest cells are induced by interactions between neural plate and adjacent tissues, and exhibit pluripotency (Buitrago-Delgado et al., 2015; Knecht and Bronner-Fraser, 2002; Pla and Monsoro-Burq, 2018; Selleck and Bronner-Fraser, 1995). Thus, the pluripotent differentiation potential of neural crest cells is ultimately derived from neural plate cells. In the most posterior region of elongating embryos, neuromesodermal progenitors generate both spinal cord and paraxial mesoderm. These progenitors are thought as derivatives of the anterior neural plate (Henrique et al., 2015; Sambasivan and Steventon, 2021).

During normal embryogenesis, neural induction in ectoderm, or restoration of neural ground state in ectodermal cells, results from inhibition of epidermalizing (or anti-neuralizing) signal, the BMP signaling, by extracellular secreted signals from the Spemann organizer (Anderson and Stern, 2016; De Robertis and Kuroda, 2004; Muñoz-Sanjuán and Brivanlou, 2002). Neural induction ensures the formation of future nervous system and also differentiation of non-neural cells. Failure of neural induction due to loss of organizer activity or the activity of embryonic neural genes causes failure in body axis formation and hence embryogenesis (De Robertis et al., 2000; Gaur et al., 2016; Rogers et al., 2009). Vice versa, neural induction by ectopic organizer activity generates formation of secondary body axis or conjoined twins. Similar effect can also be achieved by ectopic expression of genes with enriched or localized transcription in embryonic neural cells, e.g., *eed*, *yy1*, *ski*, *egfr*, *erbb2*, *erbb4*, etc. (Amaravadi et al., 1997; Nie et al., 2006; Satijn et al., 2001), which are all upregulated in cancer cells and promote cancers. In a general sense, neural induction-like process might occur in any stages and cells of an animal. Adult tissue cells could suffer various extracellular/intracellular insults, including gene mutations and dysregulations, genomic instability, microenvironmental changes, etc. These insults might occasionally cause either upregulation of neural stemness genes or genes with localized/enriched expression in embryonic neural cells or downregulation of tissue-specific genes or both, cells will gain of the property of neural stemness, i.e., the neural ground state. Consequently, the cells gain tumorigenicity and differentiation potential (Cao, 2021; Xu et al., 2021). The resulting cells proliferate quickly and meanwhile differentiate into different cell types, a process resembling embryonic growth and tissue differentiation. During normal development, neural crest cells and NSCs, both derived from neuroectoderm, migrate extensively along guided routes to differentiate further. Likewise, cancer cells migrate strongly, but without guidance. A recent single-cell RNA-sequencing analysis revealed that genes in cancer cells with a prometastatic memory predominantly relate to a neural signature (Pascual et al., 2021). To summarize, we suggest that neural stemness is a cell property unifying embryonic development and tumorigenesis (Figure 7F).

## Materials and methods

### Cell culture

HEK293T and HCT116 cells were cultured in Dulbecco’s modified eagle medium (DMEM. Thermo Fisher Scientific, #11965-092); SW480 cells was cultured in Leibovitz’s L-15 medium (L-15. Thermo Fisher Scientific, #41300039); NE-4C cells were in MEM (Gibco, #11090073) added with 1% MEM non-essential amino acids (Thermo Fisher Scientific, #11140050) and 1% Glutamax (Gibco, #35050061). All media were supplemented with 10% fetal bovine serum (FBS. Gibco, #10099141). Mouse embryonic stem cells (mESCs) were cultured in DMEM medium, added with 100 μM β-mercaptoethanol, 1 ng/ml human LIF (Cell Signaling Technology, #8911), 2 mM L-glutamine (Thermo Fisher Scientific, #25030164), 15% FBS and 1× MEM non-essential amino acids. All media were added with 50 U/ml penicillin/50 µg/ml streptomycin. For culture of NE-4C cells, petri dishes were coated with 10 µg/ml PDL (poly-D-lysine) (Sigma-Aldrich, #P0899); for mESCs, dishes were coated with 0.1% gelatin. All cells were cultured at 37°C with 5% CO_2_, except SW480, which was cultured at 37°C in 100% air. Cancer cell lines were authenticated with short tandem repeat profiling, and cells were detected free of mycoplasma contamination with PCR.

Cells were also cultured in a defined serum-free medium Ndiff^®^227 (CellArtis, #Y40002) used for derivation of primNSCs from mESCs (Ying et al., 2003) and for the test of neurosphere or neurosphere-like structure formation by NSCs or cancer cells. Control or treated cells were cultured at a density of 1×10^5^/cm^2^.

### Plasmid construction, virus packaging, cell infection or transfection

Validated MISSION^®^ short-hairpin RNAs (shRNAs) (Sigma-Aldrich) were used for knockdown of mouse Setdb1, human SETDB1 and human USP11. The shRNAs were TRCN0000092973 (mouse *Setdb1*), TRCN0000276105 (human *SETDB1*) and TRCN0000011090 (human *USP11*), which were subcloned to the lentiviral vector pLKO.1 and designated as shSetdb1, shSETDB1 and shUSP11, respectively. For stable overexpression of SETDB1, the coding region corresponding aa 110-1291 was subcloned to pLVX-IRES-Puro or pLVX-IRES-ZsGreen vector, because removal of aa 1-109 that contains nuclear export signals facilitates nuclear entry of overexpressed SETDB1 (Cho et al., 2013). The N-terminal region aa 1-725 and C-terminal region aa 726-1291 of SETDB1 were subcloned to pCS2+4×HAmcs that contains four repeats of HA-tags, and designated as HA-Nterm and HA-Cterm, respectively, and used for transient overexpression. The complete coding region for human USP11 was subcloned to pCS2+4×HAmcs vector and RPS3 was subcloned to pCS2+6×MTmcs vector that contains six repeats of Myc-tags (MT) for transient overexpression, and designated as HA-USP11 and RPS3-MT, respectively.

Virus production and cell infection were performed as described (Lei et al., 2019). Virus packaging plasmids, shRNA or overexpression constructs were co-transfected into HEK293T cells with polyethylenimine (PEI). Forty-eight hours after transfection, polybrene was added at a final concentration of 10 µg/ml to the lentiviral supernatant. The supernatant was filtered through 0.45 μm filters and centrifuged at 4°C to concentrate lentiviral particles, which were used for infecting cells. Cells after infection for 48 hours were selected with puromycin at 1 µg/ml in culture for three days if a puromycin selection vector was used. Cells were cultured further for a desired period when a significant phenotypic change was observed, or were harvested for additional assays. In parallel, virus production with the empty vector and cell infection were performed, which was used as a control for knockdown or overexpression assay, respectively.

For transient overexpression assays, HEK293T cells or HCT116 cells were transfected with an overexpression plasmid or a vector plasmid using PEI when cells grew to 70 to 80% confluency. Forty-eight hours later, cells were collected for further assays.

### Immunoblotting

Whole cell lysates were prepared for detection of protein expression using conventional SDS-PAGE and immunoblotting. Protein bands were revealed with a Western blotting substrate (Tanon, #180-501). Primary antibodies were: β-ACT (Cell Signaling Technology, #4970. 1:10,000), CCND1 (Cell Signaling Technology, #2978. 1:1,000), CDH1 (Cell Signaling Technology, #3195. 1:1,000), CDH2 (Cell Signaling Technology, #13116. 1:1,000), EIF4A1 (Abclonal, #A5294. 1:1,000), EIF4G (Cell Signaling Technology, #2469), EIF4G1 (Abclonal, #A0881. 1:1,000), EZH2 (Cell Signaling Technology, #5246. 1:2,000), FGFR1 (Cell Signaling Technology, #9740. 1:2,000), H3K27me3 (Cell Signaling Technology, #9733. 1:1,000), H3K9me3 (Abcam, #ab8898. 1:1,000), HA-tag (Cell Signaling Technology, #3724. 1:2,000), HES1 (Cell Signaling Technology, #11988. 1:2,000), LSD1 (Cell Signaling Technology, #2139. 1:2,000), MAP2 (Cell Signaling Technology, #8707. 1:1,000), MSI1 (Cell Signaling Technology, #5663. 1:1,000), MYC (Cell Signaling Technology, #13987. 1:1,000), Myc-tag (Abclonal, #AE010. 1:1,000), NEUN (Cell Signaling Technology, #12943. 1:1,000), OCT4 (Cell Signaling Technology, #83932. 1:1,000), PCNA (Cell Signaling Technology, #13110. 1:2,000), PRMT1 (Cell Signaling Technology, #2449. 1:2,000), RB1 (Cell Signaling Technology, #9309. 1:1,000), RPL24 (Abclonal, #A14255. 1:1,000), RPL26 (Abclonal, #A16680. 1:1,000), RPS3 (Abclonal, #A2533. 1:1,000), SETDB1 (Cell Signaling Technology, #2196. 1:1,000), SF3B1 (Abcam, #ab172634; Bethyl, #A300-997A-M), SOX1 (Abcam, #ab87775. 1:1,000), SOX2 (Cell Signaling Technology, #3579. 1:1,000), SOX9 (Cell Signaling Technology, #82630. 1:1,000), SRSF3 (Abcam, #ab198291. 1:1,000), SYN1 (Cell Signaling Technology, #5297. 1:1,000), TP53 (Cell Signaling Technology, #2527. 1:1,000), TUBB3 (Cell Signaling Technology, #5568. 1:1,000), USP11 (Abclonal, #A19562. 1:1,000), VIM (Cell Signaling Technology, #5741. 1:1,000), ZIC1 (Abcam, #ab134951. 1:1,000).

### Immunofluorescence

Immunofluorescence was performed as described (Lei et al., 2019). Briefly, control and treated cells were cultured in either normal medium or serum-free medium on coverslips in 6-well plates for a desired culture period. Cells were then washed with phosphate buffered saline (PBS) for three times, followed by fixation with 4% PFA for 15 min and inactivation with 50 mM ammonium chloride in PBS for 10 min. Afterwards, cells were permeabilized with 0.1% Triton X-100 for 10 min, and blocked with 0.2% fish skin gelatin (Sigma-Aldrich, #G7041) for 30 min at room temperature. Subsequently, primary antibodies were added to cells, and incubated at 4°C overnight. Primary antibodies were: EIF4G (Cell Signaling Technology, #2469. 1:200), MAP2 (Abcam, #ab183830. 1:200), MSI1 (Cell Signaling Technology, #5663. 1:200), NEFL (Cell Signaling Technology, #2837. 1:200), RPS3 (Abclonal, #A2533. 1:200), SETDB1 (Cell Signaling Technology, #2196. 1:200), SF3B1 (Abcam, #ab172634. 1:200), SOX1 (Abcam, #ab109290, 1:200), SRSF3 (Abcam, #ab198291. 1:200), SYN1(Cell Signaling Technology, #5297. 1:100), TUBB3 (Cell Signaling Technology, #5568. 1:200). Secondary antibody, Alexa Flour 594 (Thermo Fisher Scientific, #A21207, #A21203. 1:500) or anti-mouse IgG (FITC-conjugated) (Sigma-Aldrich, #F9137. 1:1000), was added to cells after washing. Cell nuclei were counterstained with DAPI. Slides were rinsed, and coverslips were mounted with an anti-fade mounting medium (Invitrogen, #S36936). Cells were viewed and photographed with a fluorescence microscope (Zeiss LSM 880).

### Reverse Transcriptase-Quantitative Polymerase Chain Reaction (RT-qPCR)

Total RNA preparation and RT-qPCR was performed as described (Yang et al., 2021). Total RNA was prepared from cells or tumors using TRIzol according to the manufacturer’s protocol. HiScript® II 1st Strand cDNA Synthesis Kit (+gDNA wiper) (Vazyme, #R212-01/02), which contains reagent for removing genomic DNA, was used for reverse transcription of cDNA. qPCR was performed on a LightCycler^®^96 System (Roche) using following parameters: one cycle of pre-denaturation at 95°C for 5 minutes, followed by 40 cycles of denaturation at 95°C for 10 seconds, annealing and extension at 60°C for 30 seconds, and an additional cycle for melting curve. In each experiment, transcription of *β-Act*/*β-ACT* was detected as a loading control. Significance in difference of transcription level was calculated based on experiments in triplicate using unpaired Student’s *t*-test. Results are presented as histograms with relative units of transcription level. Primers for qPCR are listed in Table S7.

### Cell migration/invasion assay

24-well transwell plates with inserts of 8-µm pore size (Corning, #3422) were used for cell migration assays. Each 1×10^5^ control or treated cells were suspended in 200 µl of culture medium without FBS, and added to the upper compartment. The lower compartment contained 500 µl of culture medium supplemented with 10% FBS. After incubation of the plates at 37°C for desired period as indicated in the text, cells were washed with PBS, then fixed with 37% formaldehyde, and stained with 0.5% crystal violet for 5 min. After removal of cells without migration, migrated cells were washed with PBS, and observed under a microscope and photographed.

For assessment of cell invasion, each 5×10^5^ control or treated cells were added to 80 µl of Matrigel (Corning, #354234) that was diluted in PBS at a ratio of 1:8 and evenly distributed onto a 24-well transwell insert. After incubation at 37°C for a desired period as indicated in the text, cells were processed and documented in the same way as in the migration assay.

### Soft agar colony formation assay

Soft agar was made of two layers of low melting agarose (Sangon Biotech, # A600015), with top layer being 0.35% of agarose in complete culture medium and bottom layer being 0.7% of agarose. In each well of a 6-well culture plate, 2,000 control or treated cells were plated on the top layer of agar, and cultured at 37°C for a desired period as indicated in the text. Experiments were repeated thrice. Colonies larger than 50 µm in diameter were counted for significance analysis on using unpaired Student’s *t*-test.

### Xenograft tumor assay and serial transplantation of SW480 cells

Animal use in the study was approved by and in accordance with the guidelines of the Institutional Animal Care and Use Committee (IACUC) at the Model Animal Research Center of Medical School, Nanjing University. Immunodeficient nude Foxn1^nu^ male mice of five to six week old were purchased from the National Resource Center for Mutant Mice and maintained in a specific-pathogen-free facility. Control or treated cells were suspended in 100 μl of PBS and injected subcutaneously into the dorsal flank of a mouse. Cell types and injected cell numbers are listed in Table S2. Tumor size was measured periodically before mice were sacrificed. After sacrifice of mice, tumors were excised and weighed. Tumor volume was calculated with the formula: length×width^2^/2. Significance of difference in tumor volume between control and treated groups was calculated with two-way ANOVA-Bonferroni/Dunn test. Significance of difference in tumor weight was calculated with unpaired Student’s *t*-test.

SW480 cells were passaged in vivo by serial transplantation. SW480 cells were injected subcutaneously into 6 nude mice at a does of 3×10^6^ cells per mouse. 49 days later, tumors were dissected and tumor cells were dissociated and cultured in NSC-specific Ndiff^®^227 serum-free medium for 9 days. Cell spheres formed in the serum-free medium were collected and dissociated by trituration. Each 3×10^6^ cells of the tumors at the first transplantation were injected subcutaneously into nude mice, and tumors were dissected again after 49 days after transplantation. The tumors of the second transplantation were the cell source for the third transplantation, which was done exactly as the first and second time. SW480 cells cultured in serum-free medium for 9 days (s0), sphere-forming cells from the first (s1), second (s2) and the third (s3) transplantation were subjected to assays such as RNA sequencing. Tumor volume and weight were measured, and significance in difference of tumor size and weight between different passages was analyzed in the same way as in xenograft assays above. The strategy of serial transplantation was illustrated in Figure S8A.

### Immunohistochemistry

Immunohistochemistry was used to detect expression of proteins marking different cell/tissue types in tumors using paraffin sections according to conventional method. In brief, sections were deparaffinized with a wash in xylene for 10 min first and then a wash for 5 min, and rehydrated with serial washes in 100%, 95%, 85%, 70%, 50% ethanol, and dH2O. Endogenous peroxidase activity was blocked with 3% H_2_O_2_ solution in methanol at room temperature for 15 min. After washing slides with PBS thrice, antigen retrieval was performed to unmask the antigenic epitope by treating slides with 0.01 M sodium citrate solution at 95-100°C for 20 min, followed by rinsing with PBS after cooling to room temperature. Sections were blocked with 5% BSA in PBS for 1 hour at room temperature, then incubated with primary antibody at 4°C overnight, and washed with PBS. Afterwards, biotin-conjugated goat anti-rabbit secondary antibody (Sangon Biotech, #D110066. 1:500) was added and sections were incubated at room temperature for 1 hour, and a DAB substrate (Sangon Biotech, # E670033) was added and sections were incubated at room temperature for a desired period (3-15 min) for signal visualization. Cell nuclei were counterstained with hematoxylin. Primary antibodies were: ACTA2 (Abclonal, #A11111. 1:250), AFP (Cell Signaling Technology, #4448. 1:250), BGLAP (Abclonal, #A6205. 1:500), CTSK (ABclonal, #A5871. 1:500), HES1 (Cell Signaling Technology, #11988. 1:200), KI67 (Cell Signaling Technology, #9129. 1:200), MAP2 (Abcam, #ab183830. 1:200), PAX6 (Abcam, #ab195045. 1:200), SETDB1 (Cell Signaling Technology, #2196. 1:200), SOX1 (Abcam, #ab109290. 1:200), SOX9 (Cell Signaling Technology, #82630. 1:200).

### Co-IP, mass spectrometric identification of Setdb1 interaction proteins, and functional annotation

Co-IP was performed using conventional method as described (Chen et al., 2021). Cells were collected and washed twice with ice-cold PBS. Cells were lysed by re-suspension in lysis buffer (300 mM NaCl, 1% NP-40, 2 mM EDTA, 50 mM Tris-Cl pH7.5, and protease inhibitor cocktail) on ice for 20 min. Cell lysate was centrifuged for 20 min at 12,000 rpm. Supernatant was collected and immunoprecipitation was carried out using an antibody against an endogenous protein or an HA- or MT-tag that was linked to protein G sepharose beads. In parallel, immunoprecipitation with the antibody against IgG was performed as a negative control. After incubation at 4°C overnight, beads were collected by centrifugation and washed with TBST buffer (25 mM Tris-Cl PH7.2, 150 mM NaCl, 0.5% Tween-20). Immuno-complexes were eluted by incubating the beads in 1× loading buffer at 95°C for 8 min, and then subjected to SDS-PAGE.

Identification of Setdb1 interaction proteins with mass spectrometry was performed exactly as described (Chen et al., 2021). Using the same method above, Setdb1 interaction proteins in NE-4C cells were precipitated with Setdb1 antibody; while a background control was performed in parallel with an IgG antibody. Immunoprecipitates were concentrated by SDS-PAGE, followed by coomassie blue staining. A single gel band that contained the majority of precipitated proteins was excised and subjected to in-gel digestion. Firstly, cysteine residues were reduced by addition of dithiothreitol (DTT) at a final concentration of 25 mM for 60 min at 70°C, and alkylated by addition of iodoacetamide at a final concentration of 90 mM for 30 min at room temperature in darkness. Proteins were digested with 0.2 µg of modified sequencing grade trypsin (Promega) in 50 mM ammonium bicarbonate at 37°C overnight. Peptides were extracted, dried and resupended in 10 µl of 3% acetonitrile and 2% FA, and subjected to LC-MS/MS. Peptides were analyzed by a NanoLC-2D (Eksigent Technologies) coupled with a TripleTOF 5600+ System (AB SCIEX, Concord, ON) as previously described (Zhang et al., 2020).

Data analysis was made by submitting the original files to ProteinPilot Software (version 4.5, AB Sciex). LC-MS/MS data were searched against UniProt database for *Mus musculus* (April 9, 2016, containing 50,943 sequences, http://www.uniprot.org/proteomes/UP000000589). Proteins of interests were identified by exporting mgf files from ProteinPilot, which were then subjected to Search Compare program in Protein Prospector (version 5.19.1, UCSF) for summarization, validation, and comparison of results using parameters described previously (Zhang et al., 2020). Briefly, trypsin was set as the enzyme with a maximum of two missed cleavage sites. Mass tolerance for parent ion was set at ±20 ppm, and tolerance for fragment ions was set at ±0.6 Da. The expectation value cutoff corresponding to a percent false positive (% FP) rate was determined by searching against a normal database concatenated with the reversed form of the database. Expectation values versus % FP rate were plotted by an algorithm in Search Compare. Based on the results, an expectation value cutoff corresponding to ≤0.01% FP for all peptides was chosen. At this false positive rate, false protein hits from the decoy database were not observed.

According to the result, a protein is considered to be a putative Setdb1 interaction protein when at least two peptides of a protein are identified in proteins precipitated with Setdb1 antibody but not in the precipitate with IgG antibody, or the fold change between the number of a protein peptide(s) in the sample precipitated by Setdb1 antibody and by IgG antibody is ≥ 2 (Tables S4). The proteomics data was deposited to the ProteomeXchange Consortium via the PRIDE partner repository with identifier PXD030453.

Enrichment analysis for the genes of Setdb1 binding proteins was performed using the DAVID annotation tools (Huang et al., 2007) with default settings.

### Chemical treatment of cells

PrimNSCs were derived from mESCs by culturing mESCs in serum-free Ndiff^®^227 medium at 37°C with 5% CO_2_, which formed free-floating neurospheres in the medium. To induce neuronal differentiation in primNSCs, retinoic acid (RA. Sigma-Aldrich, #R2625) was added to the medium to a final concentration of 1 μM for 6 days. For detection of protein ubiquitination and the effect of proteasomal activity on protein expression, HCT116 cells were treated with MG132 (Selleckchem, #S2619) at 25 μM for 18 hours. For detection of protein half-life, HCT116 cells infected with lentivirus carrying empty vector or SETDB1 knockdown construct, selected with puromycin, and treated with cycloheximide (CHX. Selleckchem, #S7418) at a dose of 50 μg/ml for 0, 2, 4, 6, and 8 hours, respectively. Cells were collected and subjected to additional analysis.

### Chromatin immunoprecipitation (ChIP)

HCT116 cells were infected with lentivirus derived from empty vector or SETDB1 knockdown construct. ChIP and subsequent qPCR were performed exactly as described (Lei et al., 2019). Antibodies used for ChIP were EZH2 (Cell Signaling Technology, #5246), H3K9me3 (Abcam, #ab8898), and H3K27me3 (Cell Signaling Technology, #9733). Different regions of a gene promoter were amplified from precipitated DNA using quantitative PCR. Significance in change of precipitated chromatin fragments by an antibody was calculated using unpaired Student’s *t*-test based on experiments in triplicate. Results are shown as histograms with relative units. Primers for ChIP-qPCR are listed in Table S7.

### Transcriptome profiling

Alteration of transcriptome of NE-4C cells and HCT116 cells in response to Setdb1/SETDB1 knockdown, and transcriptome of SW480 cells and cells derived from tumors from serial transplantation of SW480 cells were analyzed with RNA sequencing. Total RNA was prepared with TRIzol. Total amounts and integrity of RNA were assessed using the RNA Nano 6000 Assay Kit of the Bioanalyzer 2100 system (Agilent Technologies, CA, USA). mRNA was purified from total RNA by using poly(T) oligo-attached magnetic beads. After fragmentation of mRNA, first strand cDNA was synthesized using random hexamer primer and M-MuLV reverse transcriptase. Subsequently, RNA was degraded using RNaseH, and second strand cDNA synthesis was performed using DNA Polymerase I and dNTP. cDNA overhangs were blunted, 3’ ends of DNA fragments were adenylated. Adaptors with hairpin loop structure were ligated to prepare for hybridization. The library fragments were purified with AMPure XP system (Beckman Coulter, Beverly, USA) to select cDNA fragments of preferentially 370-420 bp in length. After qualification, cDNA library was subjected to sequencing by Illumina NovaSeq 6000. The end reading of 150bp pairing was generated. The image data measured by the sequencer were converted into sequence data (reads) by CASAVA base recognition. Raw data (raw reads) of fastq format were firstly processed through in-house perl scripts, and mapped against mouse or human reference genomes using Hisat2 (v2.0.5) software. The mapped reads of each sample were assembled by StringTie (v1.3.3b) (Mihaela Pertea.et al. 2015) in a reference-based approach. The featureCounts v1.5.0-p3 was used to count the reads numbers mapped to each gene. FPKM of each gene was calculated based on the length of the gene and reads count mapped to this gene. Differential expression analysis of two conditions was performed using the edgeR package (3.24.3). P values were adjusted using the Benjamini-Hochberg method. Padj ≤ 0.05 and |log2(fold change)| ≥ 1 were set as the threshold for significantly differential expression.

Gene Ontology (GO) and KEGG pathway enrichment analysis of differentially expressed genes was performed with the clusterProfiler R package (3.8.1). rMATS(4.0.2) software was used to analyze alternative splicing events. GATK2 (v3.8) software was used to perform SNP calling, and SnpEff (4.3q) software was used to annotate SNP. Sequencing, signal processing and data analyses were performed by Novogene Co., Ltd. (Beijing, China). RNA-seq data were deposited to the Gene Expression Omnibus (GEO) under accession numbers GSE192372, GSE192373, and GSE192374.

## Author contributions

Y.C. conceived the research. M.Z., Y.L., L.S. and L.X. performed cell, biochemical, molecular experiments and bioinformatic analysis, L.F. and M.Z. performed mass spectrometry; all authors analyzed the data; Y.C. wrote the manuscript.

