## Supplemental data for "Neural stemness unifies cell tumorigenicity and pluripotent differentiation potential"

Supplementary Figures S1-S8 and figure legends, and supplementary Tables S2 and S7 and table legends.

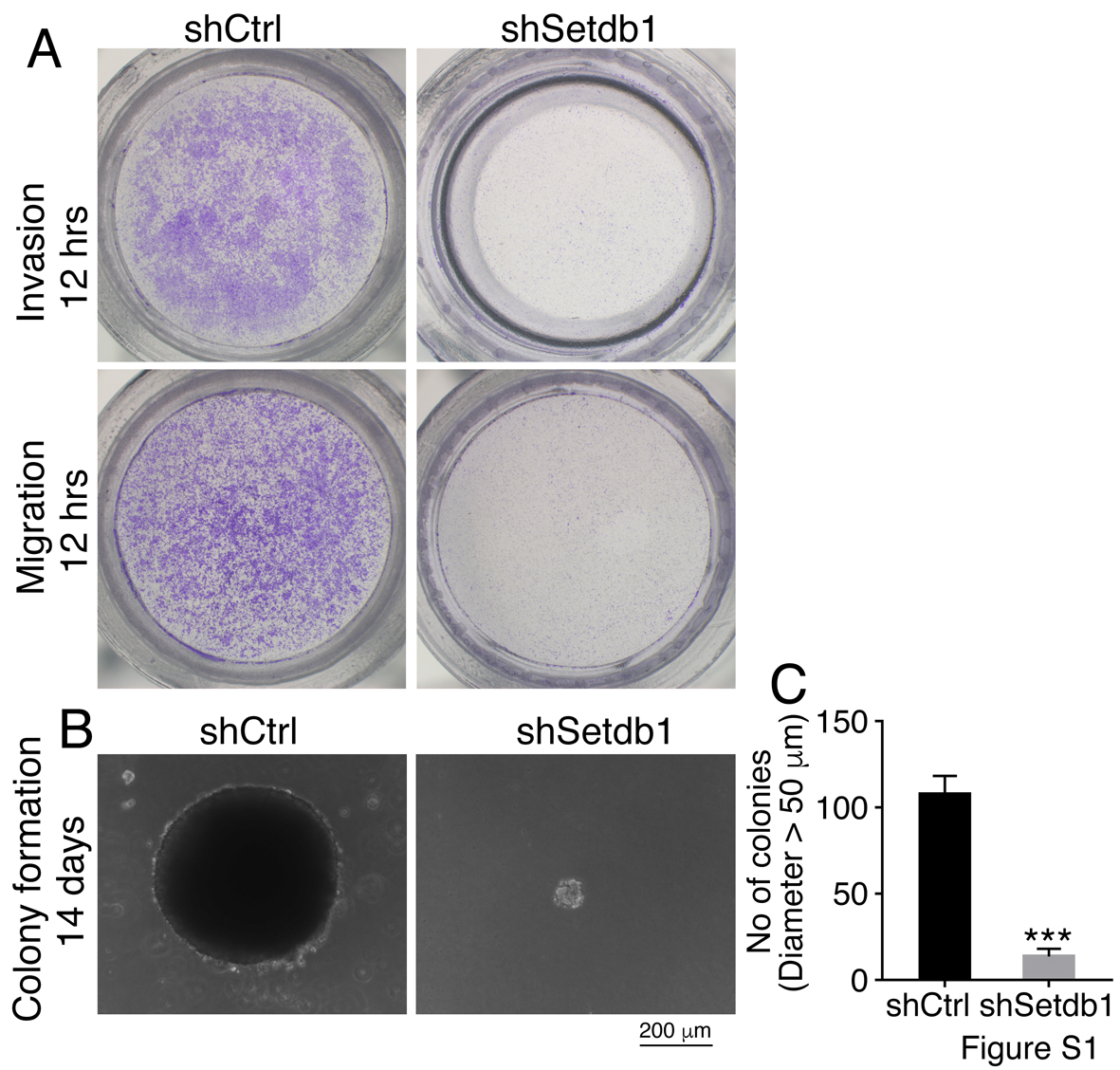

**Figure S1. Effect of knockdown of Setdb1 in NE-4C cells on cell malignant features.** (A) Change in cell invasion and migration after Setdb1 knockdown in transwell assays. (B, C) Change in colony formation in soft agar. Significance was analyzed based on experiments in triplicate using unpaired Student’s *t*-test (C). Colonies larger than 50 μm in diameter were counted. Data are shown as mean ± SEM. ***p < 0.001.

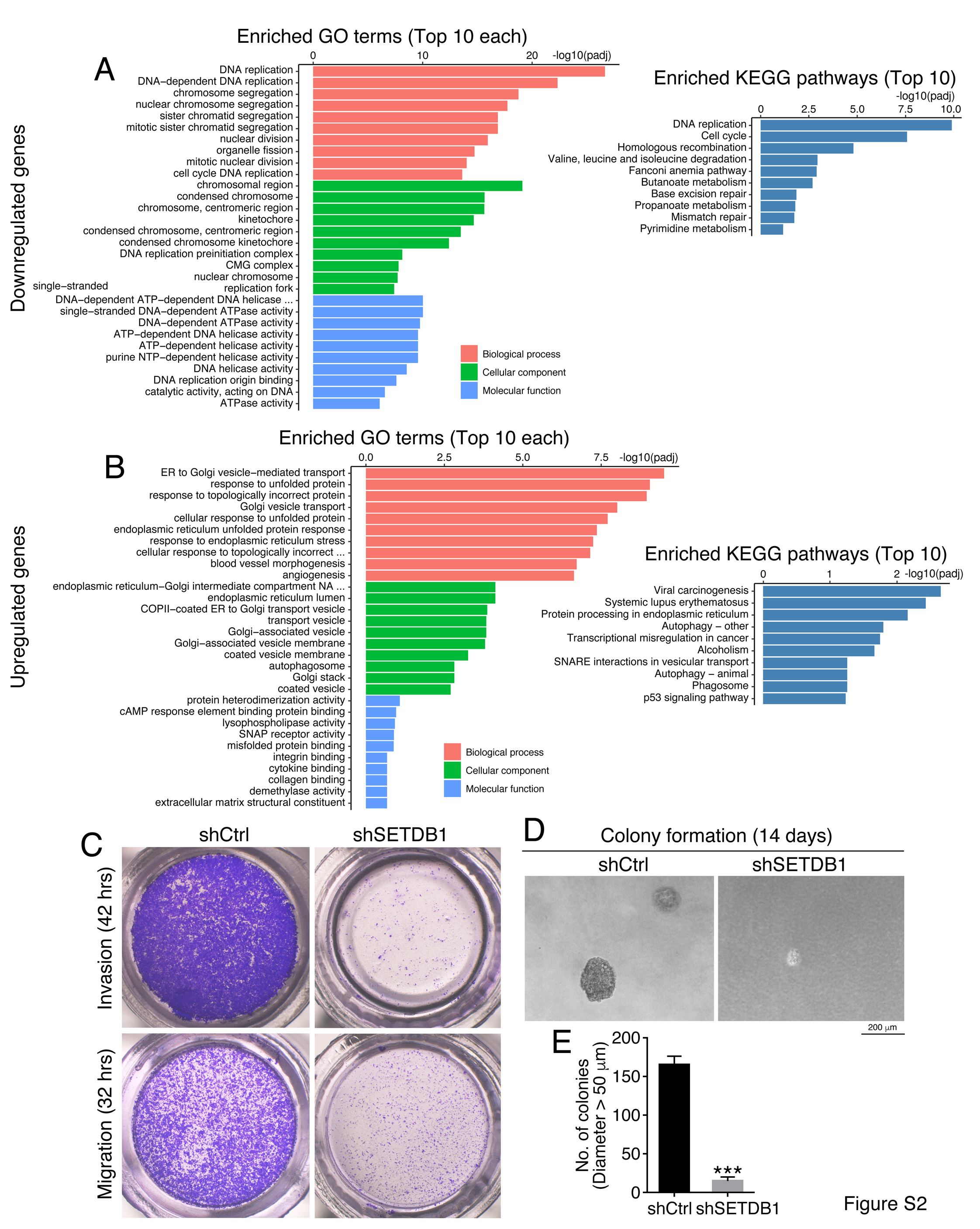

**Figure S2. Effect of SETDB1 knockdown in HCT116 cells on cell transcriptome and malignant features.** (A, B) Enrichment analysis on GO and KEGG pathway terms for downregulated (A) and upregulated (B) genes identified in a transcriptome profiling of knockdown cells. (C) Change in cell invasion and migration after SETDB1 knockdown shown by transwell assays. (D, E) Alteration in capability of colony formation in soft agar. Significance was calculated based on experiments in triplicate using unpaired Student’s *t*-test (E). Colonies larger than 50 μm in diameter were counted. Data are shown as mean ± SEM. ***p < 0.001.

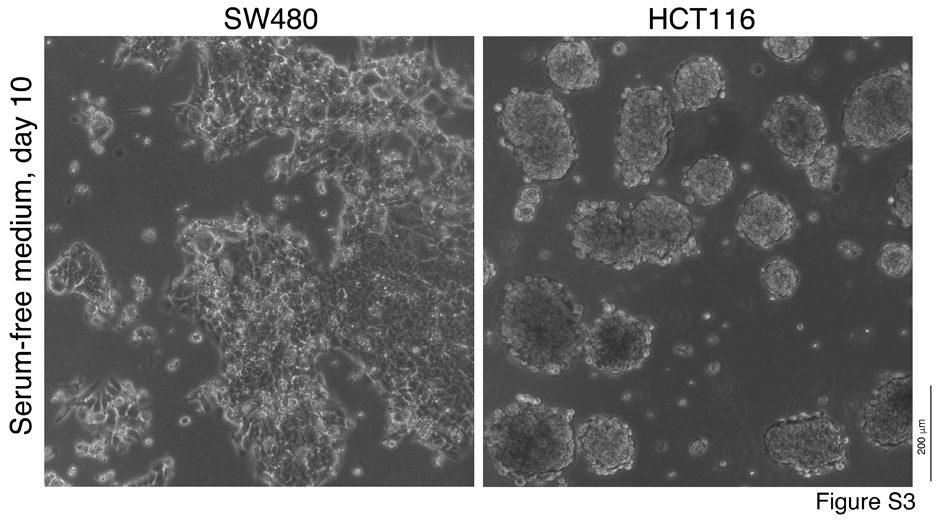

**Figure S3. The difference in capability of formation of neurosphere-like structures in NSC-specific serum-free medium by SW480 and HCT116 cells within a same period of culture.**

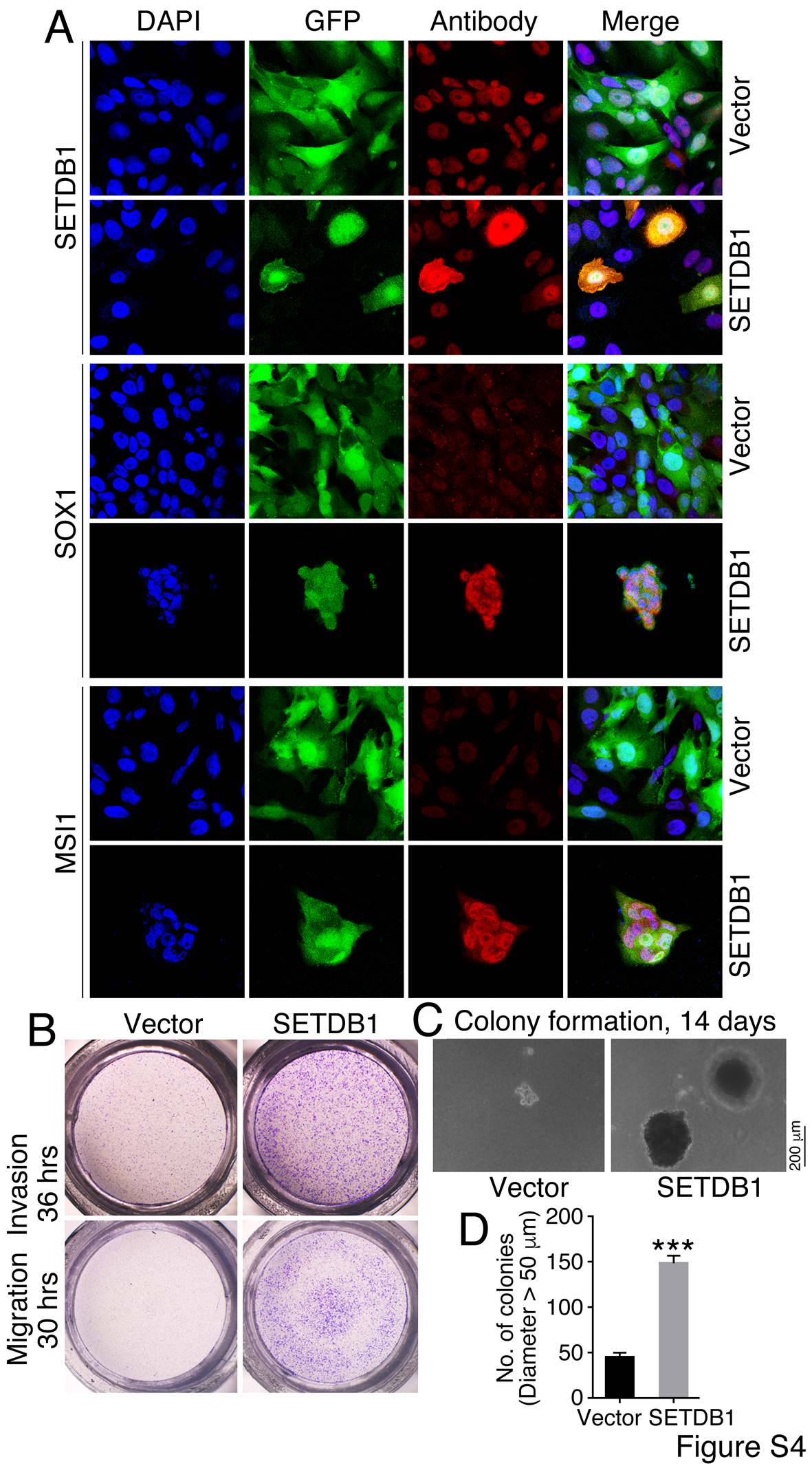

**Figure S4. Overexpression of SETDB1 in SW480 cells and the effect on cell malignant features.** (A) Effect of overexpression of SETDB1 in SW480 cells on the expression of neural stemness markers, as detected with IF. Cell nuclei were counterstained with DAPI. (B) Change in cell invasion and migration in response to SETDB1 overexpression detected with transwell assays. (C, D) Change in capability of colony formation in soft agar. Significance in change was calculated based on experiments in triplicate using unpaired Student’s *t*-test (D). Colonies larger than 50 μm in diameter were counted. Data are shown as mean ± SEM. ***p < 0.001.

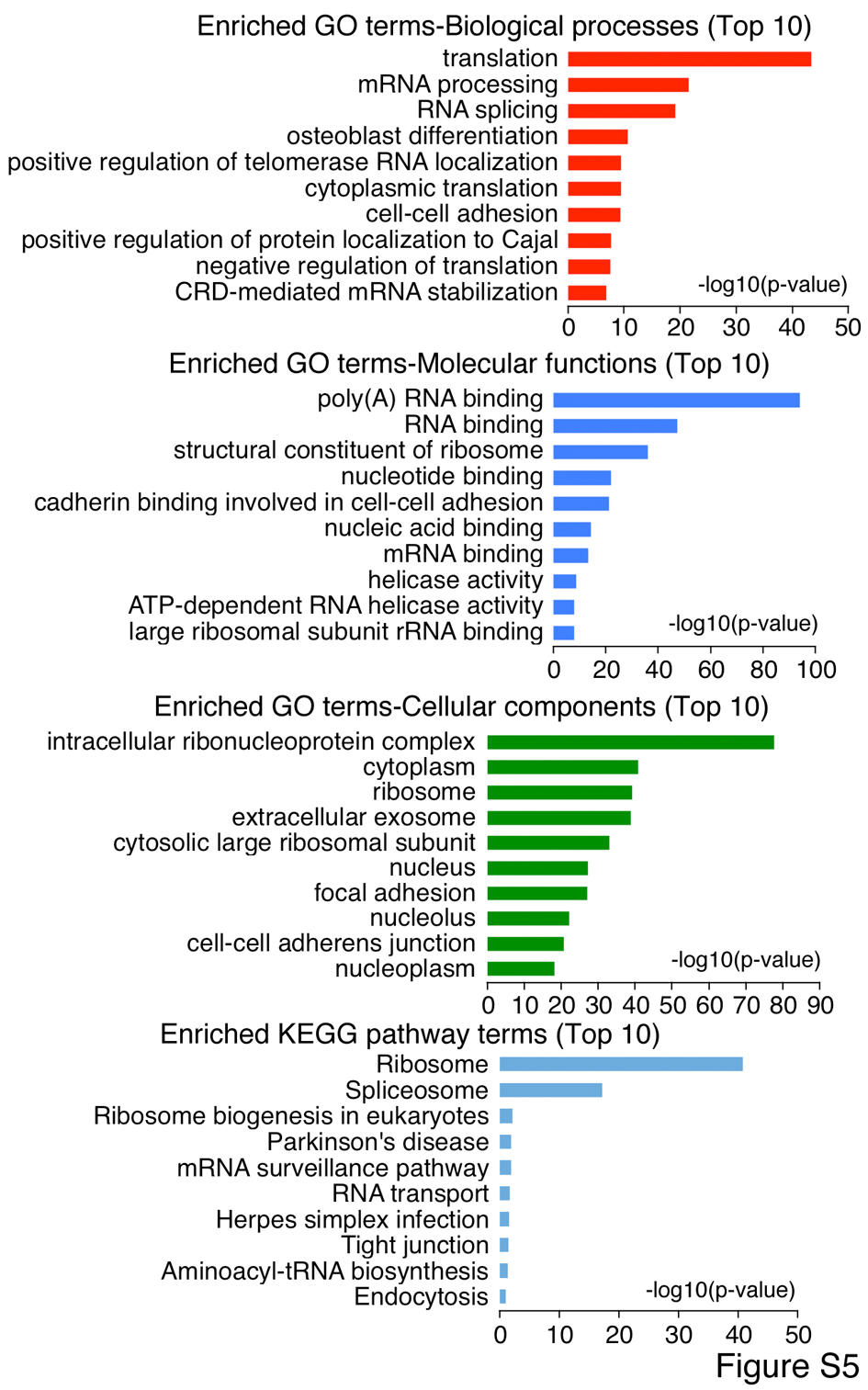

**Figure S5. Enrichment analysis on GO and KEGG pathway terms for Setdb1 interaction proteins in NE-4C cells identified by mass spectrometry.**

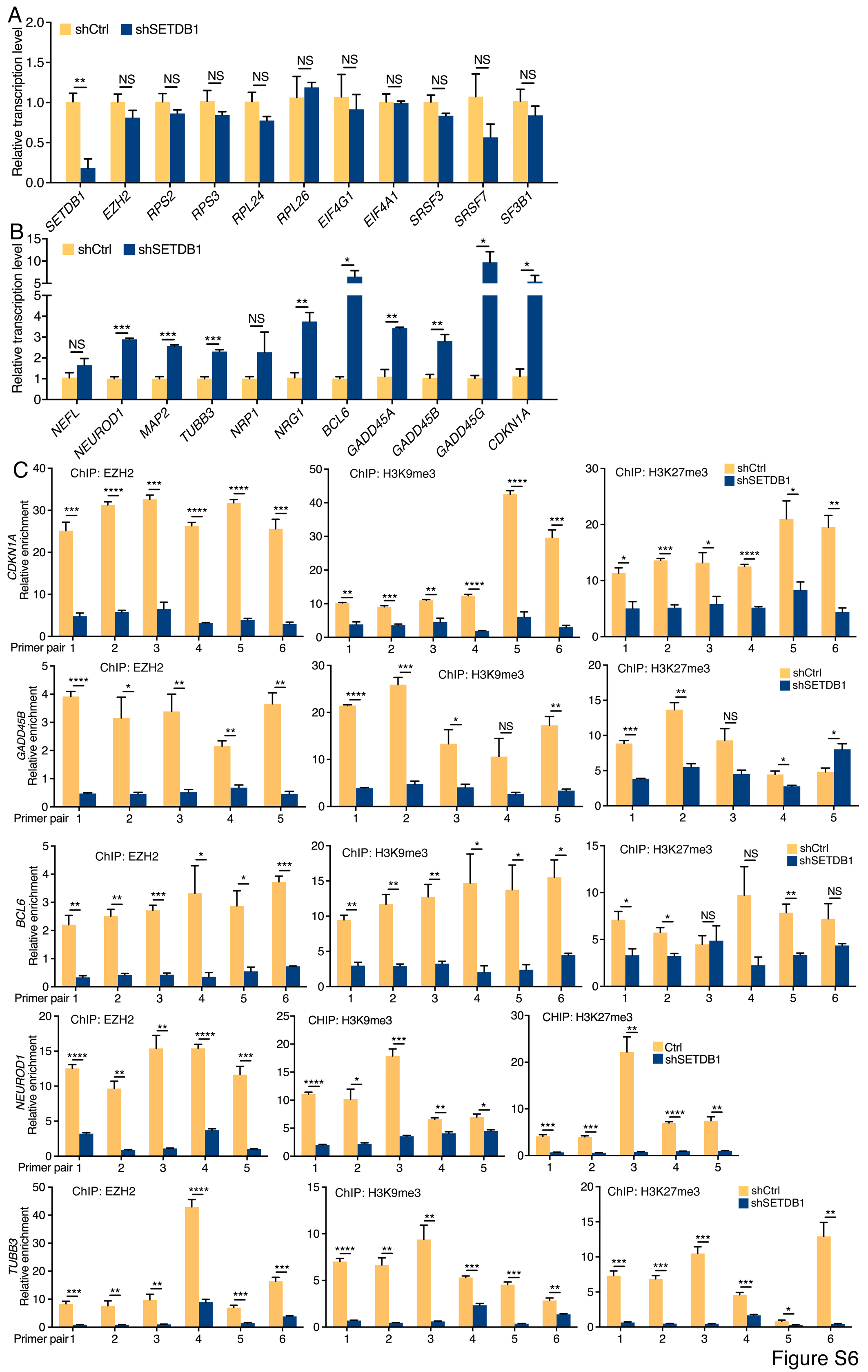

**Figure S6. Influence of SETDB1 knockdown on gene transcription and on the binding of EZH2 to promoters and the change in H3K9me3 and H3K27me3 on promoters.** (A, B) RT-qPCR detection of transcription of genes involved in epigenetic modifications, ribosome biogenesis, translation initiation and spliceosome assembly (A), and transcription of genes promoting neuronal differentiation, cell cycle and growth arrest (B) in control and knockdown HCT116 cells. Significance in transcription change was calculated based on experiments in triplicate using unpaired Student’s *t*-test. Data are shown as mean ± SEM. *p<0.05, **p < 0.01, ***p<0.001. NS: not significant. (C) ChIP detection of binding of EZH2 to promoters of *CDKN1A*, *GADD45B*, *BCL6*, *NEUROD1*, *TUBB3* and change in H3K9me3 and H3K27me3 in these promoters in response to knockdown of SETDB1 in HCT116 cells. Chromatin fragments were precipitated with antibodies against EZH2, H3K9me3, and H3K27me3, respectively, and detected with qPCR using primer pairs amplifying different regions of promoters (Table S7). Significance in difference in amplified promoter fragments was calculated using unpaired Student’s *t*-test based on experiments in triplicate. Data are shown as the mean ± SEM. *p < 0.05, **p < 0.01, ***p < 0.001, ****p < 0.0001. NS, not significant.

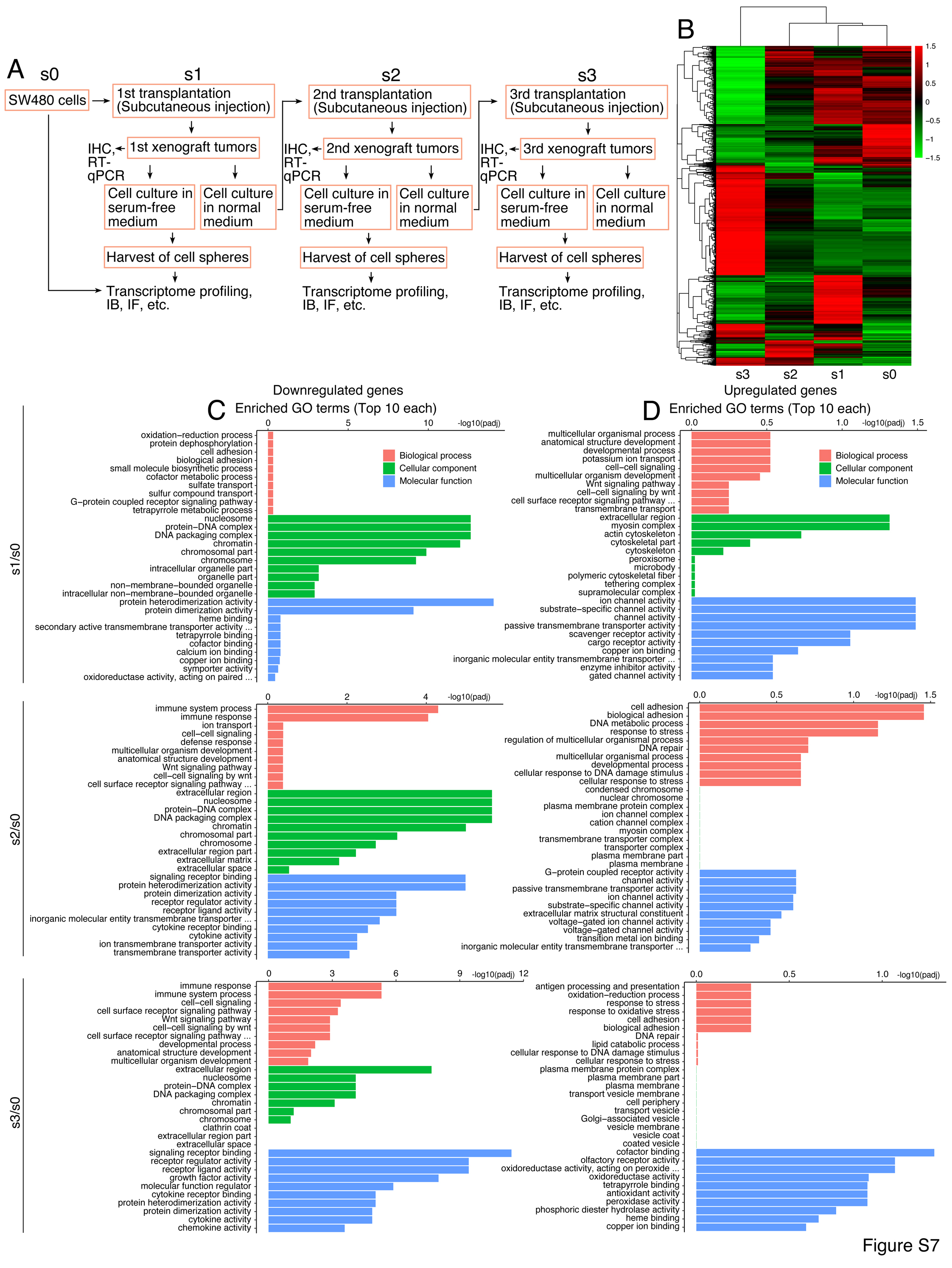

**Figure S7. Strategy of serial transplantation of SW480 cells and transcriptome profiling of cells derived from serial transplantation.** (A) Diagram depicting the strategy of serial transplantation. (B) Heatmap comparison of the transcriptomes of cells derived from s3, s2 and s1 xenograft tumors with SW480 cells cultured in serum-free medium. (C, D) Enrichment analysis on GO terms for downregulated genes (C) and upregulated genes (D) identified by transcriptome profiling in cells derived from s1, s2, s3 xenograft tumors as compared with s0 cells.

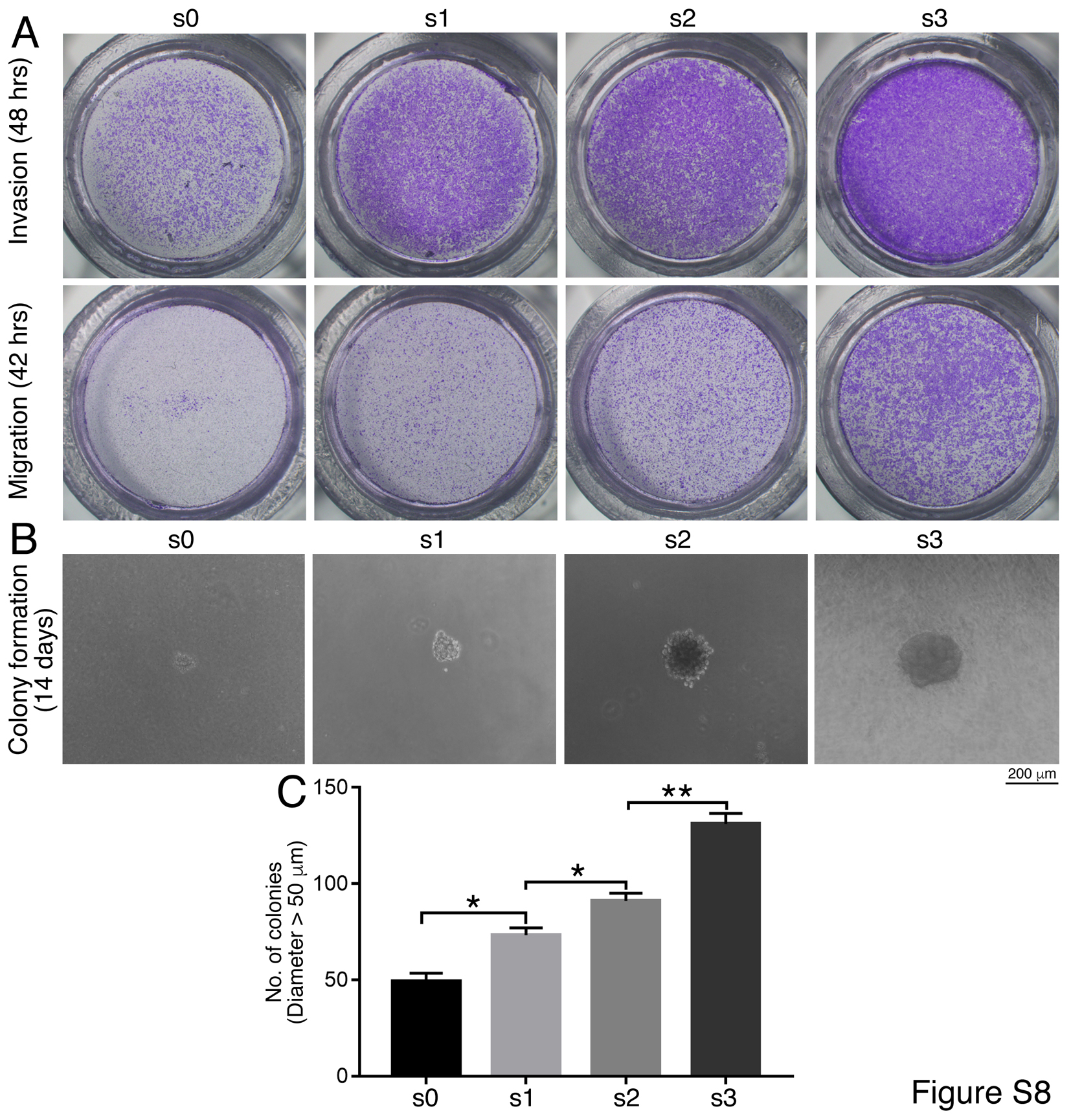

**Figure S8. Comparison of invasion, migration and colony formation by SW480 cells and cells derived from tumors of serial transplantation.** (A) Different capability of invasion and migration of cells detected with transwell assays. (B, C) Different capability of colony formation in soft agar. Significance in difference was calculated based on experiments in triplicate using unpaired Student’s *t*-test (C). Colonies larger than 50 μm in diameter were counted. Data are shown as mean ± SEM. *p < 0.05, **p < 0.01.

**Table S1. Up- and downregulated genes in NE-4C cells after knockdown of Setdb1.**

**Table S2. Xenograft analysis on different types of cells**

| Cell type | Treatment | Cells injected per mouse | Days after injection | Tumors/  Injection |
| --- | --- | --- | --- | --- |
| NE-4C | shCtrl | 1×10^6^ | 31 | 6/6 |
|  | shSetdb1 |  |  | 2/6 |
| HCT116 | shCtrl | 3×10^6^ | 28 | 6/6 |
|  | shSETDB1 |  |  | 3/6 |
| SW480 | Vector | 3×10^6^ | 42 | 6/6 |
|  | SETDB1 |  |  | 6/6 |
| SW480 (Serial transplantation) | 1^st^ injection | 3×10^6^ | 49 | 6/6 |
|  | 2^nd^ injection |  |  | 6/6 |
|  | 3^rd^ injection |  |  | 6/6 |

**Table S3. Up- and downregulated genes in HCT116 cells after knockdown of SETDB1**

**Table S4. Putative Setdb1 interaction proteins in NE-4C cells identified with mass spectrometry**

**Table S5. Alternative splicing events compared between xenograft tumor cells from high passages and lower passages during serial transplantation**

**Table S6. Numbers of SNPs in s0-s4 cells during serial transplantation**

**Table S7. Primers for RT-qPCR and ChIP-qPCR**

| Primers for RT-qPCR |  |
| --- | --- |
| Mouse genes | Primers (5’>3’) |
| *β-Act* | Forward: ccctgaagtaccccattgaa Reverse: cttttcacggttggccttag |
| *Acta2* | Forward: aatggctctgggctctgtaa  Reverse: tctcttgctctgggcttcat |
| *Ascl1* | Forward: gccaacaagaagatgagcaag Reverse: gaacccgccatagagttcaa |
| *Afp* | Forward: atgaagcaagccctgtgaac  Reverse: agcttggcacagatccttgt |
| *Cdh2* | Forward: cggtttcacttgagagcaca Reverse: catacgtcccaggctttgat |
| *Desmin* | Forward: gtgaagatggccttggatgt  Reverse: cgggtctcaatggtcttgat |
| *Foxa2* | Forward: taagcgagctaaagggagca  Reverse: gtggttgaaggcgtaatggt |
| *Gata4* | Forward: ggaagcccaagaacctgaat  Reverse: tgctgtgcccatagtgagat |
| *Klf4* | Forward: aagaggggaagaaggtcgtg  Reverse: ggtagtgcctggtcagttca |
| *Krt8* | Forward: tctgggatgcagaacatgag  Reverse: tcttcacaaccacagccttg |
| *Krt20* | Forward: tccagacttgaagcccagat  Reverse: cagccagcttagcattgtca |
| *Map2* | Forward: agcagccgaagaaacagcta  Reverse: aaggtcttgggagggaagaa |
| *Msi1* | Forward: acgtttgagagcgaggacat  Reverse: atacccagcatgaaggcatc |
| *Mturn* | Forward: agcgcaggatggacttctac  Reverse: ttcatggagggtcttcttgc |
| *Myc* | Forward: acgactccgtacagccctatt Reverse: acgtagcgaccgcaacata |
| *Myh1* | Forward: catgtccaaagccaacagtg  Reverse: acttggcgttcacagcttct |
| *Nes* | Forward: ttccctgatgatccaacctc  Reverse: acctctgtggctgcttcttt |
| *Neun* | Forward: gccatccgctacattgaga  Reverse: tgctgtcaaagctgctgttc |
| *Neurod1* | Forward: ctctggagcccttctttgaa  Reverse: tgcagggtagtgcatggtaa |
| *Oct4* | Forward: tctttccaccaggcccccgg  Reverse: cgggcggacatggggagat |
| *Pax5* | Forward: aacttgcccatcaaggtgtc  Reverse: ggcgtttgtactcagcgatt |
| *Pax6* | Forward: cacatcaggttccatgttgg Reverse: cataactccgcccattcact |
| *Pdgfra* | Forward: ctcccatccatcaaactggt  Reverse: caatctcgacgaagcctttc |
| *Robo2* | Forward: attccgttgtcaggtccaag  Reverse: acttttcccacccgattctc |
| *Setdb1* | Forward: atgtcctccctccctgggtgc Reverse: cagccgatcaacataagccac |
| *Sox1* | Forward: cacaactcggagatcagcaa Reverse: tccttcttgagcagcgtctt |
| *Sox2* | Forward: gcggagtggaaacttttgtc Reverse: tccgggaagcgtgtacttat |
| *Sox9* | Forward: gctggcaaagttgatctgaag  Reverse: gttgggtggcaagtattggt |
| *T* | Forward: ctccaacctatgcggacaat  Reverse: accattgctcacagaccaga |
| *Tubb3* | Forward: ttctggtggacttggaacct Reverse: actctttccgcacgacatct |
| *Vim* | Forward: gaccttgaacggaaagtgga Reverse: agccacgctttcatactgct |
| *Zeb2* | Forward: gggacagatcagcaccaaat Reverse: gcagtttgggcattcgtaag |
| *Zic1* | Forward: tggagccttcttccgctat Reverse: actcctcccagaagcagatgt |
| Human genes | Primers (5’>3’) |
| *ACP5* | Forward: ttccaggagacctttgagga  Reverse: tgtgggatcttgaagtgcag |
| *β-ACT* | Forward: agaaaatctggcaccacacc Reverse: tagcacagcctggatagcaa |
| *CTA2* | Forward: ctgttccagccatccttcat  Reverse: ccgtgatctccttctgcatt |
| *AFP* | Forward: agcttggtggtggatgaaac Reverse: tctgcaatgacagcctcaag |
| *BCL6* | Forward: ctgtatccagttcacccgcc  Reverse: tgaagtccaggaggatgcag |
| *BDNF* | Forward: tgttggatgaggaccagaaag  Reverse: cctcatggacatgtttgcag |
| *BGLAP* | Forward: cctttgtgtccaagcagga  Reverse: tgaaagccgatgtggtcag |
| *CDH1* | Forward: tggacagggaggattttgag Reverse: acctgaggctttggattcct |
| *CDH2* | Forward: ccatcactcggcttaatggt Reverse: acccacaatcctgtccacat |
| *CDKN1A* | Forward: cactcgtcaaatcctcccctt  Reverse: tccagtggtgtctcggtga |
| *CTSK* | Forward: tgtggtgagctttgctctgt  Reverse: gcctcaaggttatggatgga |
| *DESMIN* | Forward: tatgagaccatcgcggcta  Reverse: ggaatcgttagtgcccttca |
| *EIF4A1* | Forward: atgtctgcgagccaggat  Reverse: cagtcccagattgggctt |
| *EIF4G1* | Forward: atgaacaaagctccacag  Reverse: ggctgcactctgcactcg |
| *EZH2* | Forward: agggcacagcagaagaactaa  Reverse: cgcctacagaaaagcgtatga |
| *FOXA2* | Forward: cccacaaaatggacctcaag  Reverse: ataatgggccgggagtaca |
| *GADD45A* | Forward: atgactttggaggaattctcg  Reverse: cagcgtcggtctccaagag |
| *GADD45B* | Forward: atgacgctggaagagctcg  Reverse: gactggatgagcgtgaagtg |
| *GADD45G* | Forward: ctggaagaagtccgcggc  Reverse: gcaggtcgcccggcgcac |
| *GATA6* | Forward: gtgtgcaatgcttgtggact  Reverse: agttggagtcatgggaatgg |
| *HNF1A* | Forward: cctcaaagagctggagaacct Reverse: ttgttgaggtgttgggacag |
| *HNF4A* | Forward: gtgtccatacgcatccttga Reverse: tactggcggtcgttgatgta |
| *KRT20* | Forward: atgaagtcatggcccagaag  Reverse: ttcatgctgagatgggactg |
| *MAP2* | Forward: cagggaggaatttgtggaga  Reverse: atggtctcctttccacctca |
| *MSI1* | Forward: accaagagatccaggggttt Reverse: tcgttcgagtcaccatcttg |
| *MTURN* | Forward: acgcaggatggatttctacg  Reverse: acttcgtggagcgtcttctt |
| *NEFL* | Forward: agaccctggaaatcgaagca  Reverse: tcgccttccaagagtttcct |
| *NEUROD1* | Forward: gacgatcaaaagcccaagag  Reverse: tggacagcttctgcgtctta |
| *NRG1* | Forward: gagttggcaccacagccttg  Reverse: tccagaatcagccagtgatg |
| *NRP1* | Forward: tgccgctcctctgcgccg  Reverse: ccaaatcgaagtgagggttg |
| *OCT4* | Forward: tgagaggcaacctggagaat  Reverse: cagcagcctcaaaatcctct |
| *PAX6* | Forward: aagggccaaatggagaagag  Reverse: gccagatgtgaaggaggaaa |
| *PDGFRA* | Forward: ggccccatttacatcatcac  Reverse: catagctccgtgtgctttca |
| *MYC* | Forward: tcaagaggcgaacacacaac Reverse: atgagcttttgctcctctgc |
| *RPL24* | Forward: atgcgagtcggctttccttt Reverse: agagatgcaccagtaatggcc |
| *RPL26* | Forward: tacgtgatctacatcgagcgc Reverse: tctcctccttgtacttgccct |
| *RPS2* | Forward: tgaagatcaagtccctggagg Reverse: aaatgccttgaacctggtgc |
| *RPS3* | Forward: atgctgaaaaggtggccact Reverse: tttcccagacaccacaacct |
| *SETDB1* | Forward: gtggatggcagcctagtc  Reverse: tggcttgaactgggttcc |
| *SF3B1* | Forward: atggcgaagatcgccaag  Reverse: tctgaccaagcaaactcg |
| *SOX1* | Forward: aagtcaaaacgaggcgagag  Reverse: aagtgcttggacctgcctta |
| *SOX2* | Forward: catcacccacagcaaatgac Reverse: cctccccaggttttctctgta |
| *SOX9* | Forward: ccgaagaaagagaggaccaa  Reverse: gcgcttggataggtcatgtt |
| *SRSF3* | Forward: atgcatcgtgattcctgt  Reverse: cattcgacagttccactc |
| *SRSF7* | Forward: atgtcgcgttacgggcgg  atccagtcctcgtactgc |
| *SYNJ1* | Forward: tggcttctgaacagttggtg  Forward: agcaaaggctggttgtatgg |
| *T* | Forward: aagtacgtgaacggggaatg  Reverse: tgagcttgttggtgagcttg |
| *TUBB3* | Forward: gtgcggaaggagtgtgaaa  Reverse: acgacgctgaaggtgttcat |
| *ZEB2* | Forward: caccaaatgctaacccaagg  Reverse: tgtgcgaactgtaggaacca |
| *ZIC1* | Forward: acaaaaggacgcacacagg  Reverse: ggattcgtggaccttcatgt |
| *ZIC2* | Forward: gtacggccccatgaatatga  Reverse: cttgggattgctcagttgct |
| Primers for ChIP-qPCR |  |
| Gene | Primers (5’>3’. Predicted amplified regions are indicated by numbers at the end of each reverse primer. The first base of the transcriptional start site is designated as +1.) |
| *BCL6* | Primer pair 1  Forward: cgcactccccctcttatgtc  Reverse: tttggaggttccggttcgag(-380/-184)  primer pair 2  forward: tgaacccaccgaaaactgct  Reverse: cacaaaagggagcgagaggt (-621/-451)  Primer pair 3  Forward: gattcgtgcggctgtgtttt  Reverse: tccaaatctcggttcggctc(+606/+850)  Primer pair 4  Forward: tggttcaaacctctcgctcc  Reverse: aagacgatggtatggcctgc(-478/-243)  Primer pair 5  gtggctttgagggcttttgg  ataatcacctggtgtccggc(-127/+71)  Primer pair 6  tgccgaagattagtcccacg  cagactagcccgaatcaccc(+285/+515) |
| *CDKN1A* | Primer pair 1  Forward: cttcaaggcagtgggagaag  Reverse: gattgtggctaaaccccaga(+553/+711)  Primer pair 2  Forward: aggaaggggatggtaggaga  Reverse: ctcccagcacacactcacac (+39/+188)  Primer pair 3  Forward: gaggcagaattgcttgaacc  Reverse: ataggggcagtcagctttca (-566/-339)  Primer pair 4  Forward: tctcagctcactgcaacctc  Reverse: tggtggcttacgcctgtaat (-985/-761)  Primer pair 5  Forward: gcaggtgtgatgaccaacaa  Reverse: tttcatccattcattcaaaaacc(-1331/-1139)  Primer pair 6  Forward: ctgaggggaggctcatactg  Reverse: agagaggcatcctccagaca (-1884/-1475) |
| *GADD45B* | Primer pair 1  Forward: tgtgtgagtcagaccccctt  Reverse: cgatccacaaaacggcagag (+502/+657)  Primer pair 2  Forward: cgcagaagtaagtagccggg  Reverse: gggggtctgactcacacatt (+270/+519)  Primer pair 3  Forward: tgcactcgcccttgtctt  Reverse: cgggaagcagcgaaatcc (-104/+113)  Primer pair 4  Forward: gctggaaatcccgcgcgc  Reverse: ctctgcggttggccgacg (-307/-110)  Primer pair 5  Forward: gaaggtcttggacgagcg  Reverse: gttgtcgcacgccacgagc (+113/+265) |
| *NEUROD1* | Primer pair 1  Forward: ttttacgcacattgggagtg  Reverse: agcggtaacaggtagcagga (+627/+793)  Primer pair 2  Forward: aggccactcgctctgatcta  Reverse: cctttgtggccagaagaaag (-367/-184)  Primer pair 3  Forward: agctcgctttgaggacaaga  Reverse: atcgtcctctcccagttcct (-843/-676)  Primer pair 4  Forward: gagttcgcagccattaatcc  Reverse: tagtccaggcattgaccaca (-1248/-1034)  Primer pair 5  Forward: ctggctaggaccctcttcct  Reverse: agacctgaaaccctctgcaa (-1669/-1490) |
| *TUBB3* | Primer pair 1  Forward: ccgacgctttgtttcttctc  Reverse: ggctttgtacggagggtctt (+802/+975)  Primer pair 2  Forward: ggggttcgtctgtacatcgt  Reverse: tcagagaaggaagggagcaa (+291/+450)  Primer pair 3  Forward: tctcgctgaagagaccacct  Reverse: aggaagcagctcccagttct (-318/-138)  Primer pair 4  Forward: aaaccccagcctagaggaag  Reverse: caggcatctcagagtggaca (-811/-595)  Primer pair 5  Forward: cctagtggtatggggctgag  Reverse: tttctggaaggggctaaggt (-1224/-1017)  Primer pair 6  Forward: ctccaggcaggacttctcac  Reverse: gatgctgctcagtcattgga (-1607/-1444) |
